# CXCL8/CXCR2 signaling mediates bone marrow fibrosis and represents a therapeutic target in myelofibrosis

**DOI:** 10.1101/2021.12.08.471791

**Authors:** Andrew Dunbar, Dongjoo Kim, Min Lu, Mirko Farina, Julie L. Yang, Young Park, Francesca Gobbo, Paola Verachi, Fabrizio Martelli, Abdul Karzai, Wenbin Xiao, Lijuan Xia, Nada Elmansy, Maria Kleppe, Zhuo Chen, Yang Xiao, Erin McGovern, Jenna Snyder, Aishwarya Krishnan, Corrine Hill, Keith Cordner, Anouar Zouak, Mohamed E. Salama, Jayden Yohai, Eric Tucker, Jonathan Chen, Jing Zhou, Tim McConnell, Richard Koche, Raajit Rampal, Anna Rita Migliaccio, Rong Fan, Ross L. Levine, Ronald Hoffman

**Affiliations:** Human Oncology & Pathogenesis Program, Memorial Sloan Kettering Cancer Center, New York, NY, USA; Leukemia Service, Department of Medicine and Center for Hematologic Malignancies, Memorial Sloan Kettering Cancer Center, New York, NY, USA; Myeloproliferative Neoplasm-Research Consortium; Department of Biomedical Engineering, Yale University, New Haven, CT, USA; Division of Hematology/Oncology, Tisch Cancer Institute and Department of Medicine, Icahn School of Medicine at Mount Sinai, New York, NY, USA; Blood Diseases and Bone Marrow Transplantation Unit, Cell Therapies and Hematology Research Program, Department of Clinical and Experimental Sciences, University of Brescia, Brescia, Italy; Center for Epigenetics Research, Memorial Sloan Kettering Cancer Center, New York, NY, USA; Department of Veterinary Medical Sciences, University of Bologna, Italy; Department of Biomedical and Neuromotor Sciences, University of Bologna, Italy; Istituto Superiore di Sanità (ISS), Rome, Italy; Department of Pathology, Memorial Sloan Kettering Cancer Center, New York, NY, USA; Department of Pathology, Mayo Clinic School of Medicine, Rochester, MN; IsoPlexis Corporation, Branford, CT, USA; Altius Institute for Biomedical Sciences, Seattle, WA, USA; Università Campus Bio-Medico di Roma, Rome, Italy

**Keywords:** Myeloproliferative neoplasms, myelofibrosis, pro-inflammatory signaling, IL-8, CXCL8/CXCR2

## Abstract

Pro-inflammatory signaling is a hallmark feature of human cancer, including in myeloproliferative neoplasms (MPNs), most notably myelofibrosis (MF). Dysregulated inflammatory signaling contributes to fibrotic progression in MF; however, the individual cytokine mediators elicited by malignant MPN cells to promote collagen-producing fibrosis and disease evolution remain yet to be fully elucidated. Previously we identified a critical role for combined constitutive JAK/STAT and aberrant NF-κB pro-inflammatory signaling in myelofibrosis development. Using single-cell transcriptional and cytokine-secretion studies of primary MF patient cells and two separate murine models of myelofibrosis, we extend this previous work and delineate the role of CXCL8/CXCR2 signaling in MF pathogenesis and bone marrow fibrosis progression. MF patient hematopoietic stem/progenitor cells are enriched in a CXCL8/CXCR2 gene signature and display dose-dependent proliferation and fitness in response to exogenous CXCL8 ligand *in vitro*. Genetic deletion of *Cxcr2* in the hMPL^W515L^ adoptive transfer model abrogates fibrosis and extends overall survival, and pharmacologic inhibition of the CXCR1/2 pathway improves hematologic parameters, attenuates bone marrow fibrosis, and synergizes with JAK inhibitor therapy. Our mechanistic insights provide a rationale for therapeutic targeting of the CXCL8/CXCR2 pathway in MF patients at risk for continued fibrotic progression.

## INTRODUCTION

Primary and secondary forms of myelofibrosis (MF) are clonal myeloproliferative neoplasms (MPNs) characterized by frequent constitutional symptoms, progressive cytopenias, splenomegaly, increased bone marrow microvessel density, megakaryocytic hyperplasia and atypia, and increased risk of evolution to acute leukemia (often termed blast phase myeloproliferative neoplasm (MPN-BP)).^1^ Overt MF can evolve from other forms of MPN, which include polycythemia vera (PV) and essential thrombocythemia (ET). Gain-of-function mutations of the JAK/STAT pathway are present in the great majority of MPN patients highlighting the role of constitutive JAK/STAT activation in disease initiation and maintenance.^2–7^ While the recurrent mutations in MPNs have been extensively studied,^8^ the phenotypic and prognostic pleiotropy observed across MPN sub-types, including MF, suggests that biologic factors in addition to activated JAK/STAT signaling contribute to MF progression and eventual MPN-BP transformation.

Chronic aberrant pro-inflammatory cytokine signaling is an important mediator of fibrotic development across multiple tissue types, including the bone marrow.^9^ Bone marrow fibrosis can occur as a consequence of several different autoimmune and cytokine-mediated conditions underscoring the contribution of inflammatory signaling on fibrotic progression independent of constitutive JAK/STAT activation.^10^ Recent single-cell studies and others have provided insight into how chronic pro-inflammatory changes within the bone marrow compartment promote mesenchymal stromal cell (MSC) remodeling to drive fibrosis.^11–13^ In addition, bone marrow derived fibrocytes are thought to represent an alternative source for myofibroblast proliferation during skin wound healing and in liver, lung, and kidney fibrosis, as well as in the stromal reaction to MF.^14,15^ Importantly, many of the same pathways shown to contribute to myofibroblast differentiation are also frequently implicated in MPN hematopoietic stem cell (HSC) expansion/differentiation, including TGFβ,^16–18^ JAK/STAT,^19^ and TNFα^20,21^ highlighting an important cross-talk between clonal MPN HSCs and stromal cells within the bone marrow niche.^22,23^ Previously, we and others have shown both JAK/STAT and TNFα/NF-κB inflammatory pathways cooperate to promote marrow fibrosis in MPNs.^24,25^ Canonical NF-κB activation elicits a myriad of chemokines and cytokines that contribute to the acute and chronic inflammatory response, including IL6, CXCL8 (IL-8), and MIP-1α, among others.^26^ These cytokines, in addition to other pro-fibrogenic cytokines (e.g. TGFβ) are frequently elevated in patients across the spectrum of MPNs, are enriched in MF, and have prognostic relevance.^27,28^ Recent studies highlight the contribution of specific cytokines in MF progression.^29^ Additional cytokine pathways are likely involved however, and the full spectrum of individual pro-inflammatory cytokines which play causative roles in MF have yet to be identified. Furthermore, the contribution of pro-inflammatory pathways might not be static as the disease evolves. In a recent study, NF-κB was enriched in early fibrotic development but down-regulated in more advanced stages of fibrosis suggesting clonal HSCs might elicit adaptive and evolving changes on surrounding stroma over time.^29^

We hypothesized that there are specific cytokines/chemokines elicited by MPN HSCs which promote MF development and might predict the likelihood of disease progression. Using both single-cell transcriptional and cytokine platforms, we identify increased CXCL8/CXCR2 signaling in MF compared to other MPN subtypes and assess the role of CXCL8/CXCR2 signaling in MF pathogenesis and therapeutic response.

## RESULTS

### CXCL8/CXCR2 signaling is enriched in CD34+ hematopoietic stem/progenitor cell populations (HSPCs) of fibrotic MPN

To explore specific cytokine pathways in MPN HSCs with enriched NF-κB activity (and thus fibrosis), we carried out single-cell gene expression profiling (scRNA-seq) on isolated CD34+ cells from 7 separate individual MPN patients across MPN sub-types with varying degrees of fibrosis **(Supp. Table S1)** using an enclosed microwell array platform^30^ with a focus on NF-κB pathway-associated cytokine mediators. Cell populations were detected across multiple hematopoietic cell lineages **(Supp. Figure 1A)**, with overall enrichment in CD34+ HSPC populations consistent with our CD34+ positive selection **(Supp. Figure 1B-C)**. The most predominant cell population detected across patients was the common myeloid progenitor (CMP) population, and the lineage distribution across patients was mostly similar **(Supp. Figure 1D)**.

Following filtration, a total of 97,412 genes in 8,144 single cells were included for final analysis of differentially expressed genes. Uniform manifold approximation and projection (UMAP) revealed tight segregation of mapped HSCs cells on a per-patient basis, with significant clustering of patients ET3 and MF1 separate from the other patients evaluated (**Figure 1A**). Similar segregation was also observed when comparing other cell fractions from these patients, including both CMPs and granulocyte-monocyte progenitors (GMPs) **(Supp. Figure 1E)** suggesting highly shared transcriptional programs between these two individuals across cell types. Notably, these patients had the greatest degree of bone marrow fibrosis as evidenced by bone marrow reticulin stains assessed at the time of sample collection suggesting enrichment in a “fibrotic” gene signature independent of clinical MPN sub-type **(Supp. Table S1;** **Figure 1B**). Unsupervised hierarchical clustering and gene ontology analysis of these 2 patients revealed enrichment in pathways recently implicated in MF, including primarily TNFα signaling via NF-κB and inflammatory response pathways (**Figure 1C-D**). Of the most enriched specific cytokine/chemokine pathways known to be regulated by TNFα/NF-κB in “high fibrosis” patients, we observed a signature suggestive of CXCR2-related chemokine activation, including overexpression of the cytokine/chemokines *CXCL2* (MIP-2α), *CXCL3* (MIP-2β), and *CXCL8* (IL-8), all of which signal through the CXCR1/2 receptors.^31^ Similar enrichment in chemokine signaling and up-regulation of *CXCL2*, *CXCL3*, and *CXCL8* in these patients was also observed in CMPs **(Supp. Figure 1F)** suggesting a shared transcriptional program enriched in CXCR2 signaling into myeloid lineage commitment. Intriguingly, upregulation of the lncRNAs MALAT1 and NEAT1 was also observed in the “high fibrotic” cell clusters. MALAT1 has an emerging role as a potent regulator of pro-inflammatory signaling and was recently shown to directly elicit *CXCL8* expression through epigenetic modulation of the *CXCL8* promoter via p300.^32,33^ Together, these data suggested that the CXCL8-CXCR2 axis is enriched in CD34+ HSPCs of fibrotic MPN patients.

**Figure 1:**
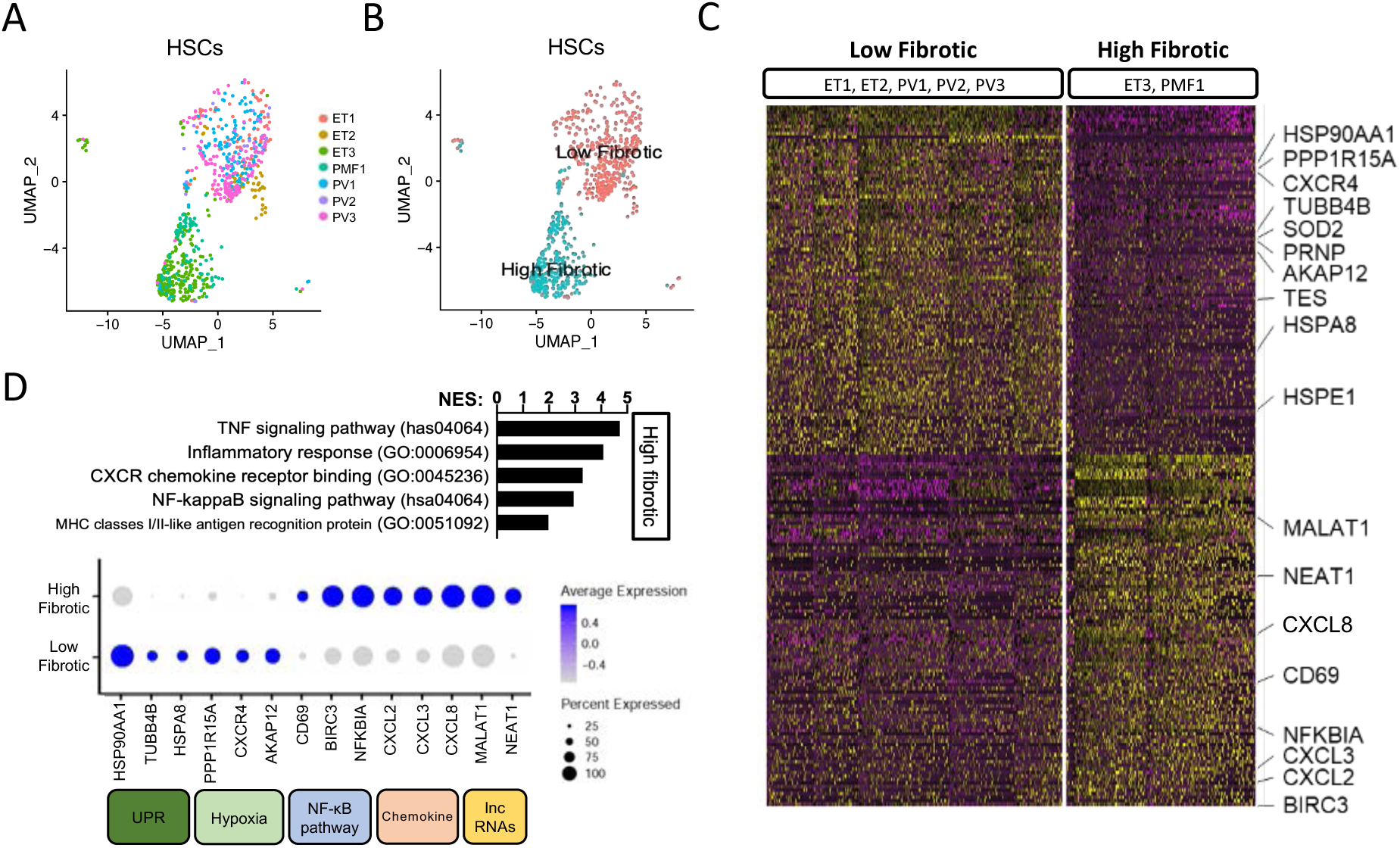
Single-cell transcriptional profiling (scRNA-Seq) identifies genes and gene sets enriched in MF, including the chemokines CXCL1, CXCL2, and CXCL8 and their receptor CXCR2. **(A)** Uniform manifold approximation and projection (UMAP) visualization of individual hematopoietic stem cells (HSCs) colored by patient (ET, essential thrombocythemia; PV, polycythemia vera; PMF, primary myelofibrosis; See Supp. Table S1). **(B)** UMAP clustering of HSCs colored by “low” vs. “high” fibrosis patients. **(C)** Hierarchical clustering of differentially expressed genes in “low fibrotic” and “high fibrotic” MPN patients. Adjusted p-value <0.01 (Wald test) and abs (log2-fold change) >1. Purple, negative values; yellow, positive values. N=2-5/group. **(D)** Top Panel: Gene set enrichment analysis (GSEA) of differentially expressed genes of high vs. low fibrotic patient HSCs (NES, normalized enrichment score); lower panel: most differentially expressed genes and their pathway associations (pct exp: % of cells expressing listed gene; avg. exp. scale: Z score of normalized read counts, with blue, positive values and gray, negative values).

### Single-cell cytokine arrays identify enrichment in CXCL8 secreting CD34+ cells in a subset of MF patients

Previous studies have highlighted the prognostic relevance of serum CXCL8 levels in MF.^27^ In order to identify individual cytokines secreted by MF HSPCs, we utilized a microchip system^34^ allowing for the detection of single cell cytokine secretion from CD34+ cells isolated from clinical samples across MPN sub-types (N=11 MF, N=13 PV, N=14 ET; **Supp Table S2)**. To allow for an expanded patient cohort, we limited the number of cytokines assayed to a few of those known to be mediated through NF-κB, specifically CXCL8, MIP-1β, TNFα, IL-6, and RANTES. Of these 5 cytokines assessed, CXCL8-only secreting cells were highly enriched in MF patients as compared to PV and ET patients (54% MF vs. 31% PV vs. 0% ET) (**Figure 2A**). Consistent with our previous study,^35^ the majority of detectable cells secreted only one cytokine **(Supp. Figure 2A)**, and the number and complexity of cytokines enumerated from individual cells (poly-functionality index) did not vary significantly across MPN sub-types **(Supp. Figure 2B)**. Intriguingly, RANTES was significantly enriched in ET patients **(Supp. Figure 2C)**, but given the transcriptional data highlighting CXCL8 in MF and previous associations of CXCL8 on clinical outcome in this MPN sub-type, we decided to focus our analysis on CXCL8 specifically. The frequency of CXCL8 secreting CD34+ cells not only correlated with MF disease phenotype (**Figure 2B**), but also with the degree of reticulin fibrosis and leukocytosis **(Figure 2C, Supp. Figure 2D)**, suggesting that the extent of CXCL8 secretion may be associated with fibrotic progression and/or serve as a biomarker for the presence of significant bone marrow fibrosis (**Figure 2D**). General enrichment of CXCL8 levels in MF was further supported by cytokine profiling of CXCL8 levels in collected plasma of MPN patients (**Figure 2E**). To assess CXCL8 expression in situ, we performed IHC for CXCL8 on sectioned whole patient bone marrow in patients with MF and observed increased CXCL8 expression in megakaryocytes and splenic stromal cells in a subset of MF patients (8/15) not observed in healthy controls (0/4) (**Figure 2F**). Consistent with this, we identified increased CXCL8 ligand secretion from cultured MF megakaryocytes (MKs) in liquid culture in addition to the known pro-fibrogenic cytokines VEGF and TGFβ **(Supp. Figure 3A)**. This was also confirmed at the transcriptional level by quantitative PCR of isolated cells **(Supp. Figure 3B)**.

**Figure 2:**
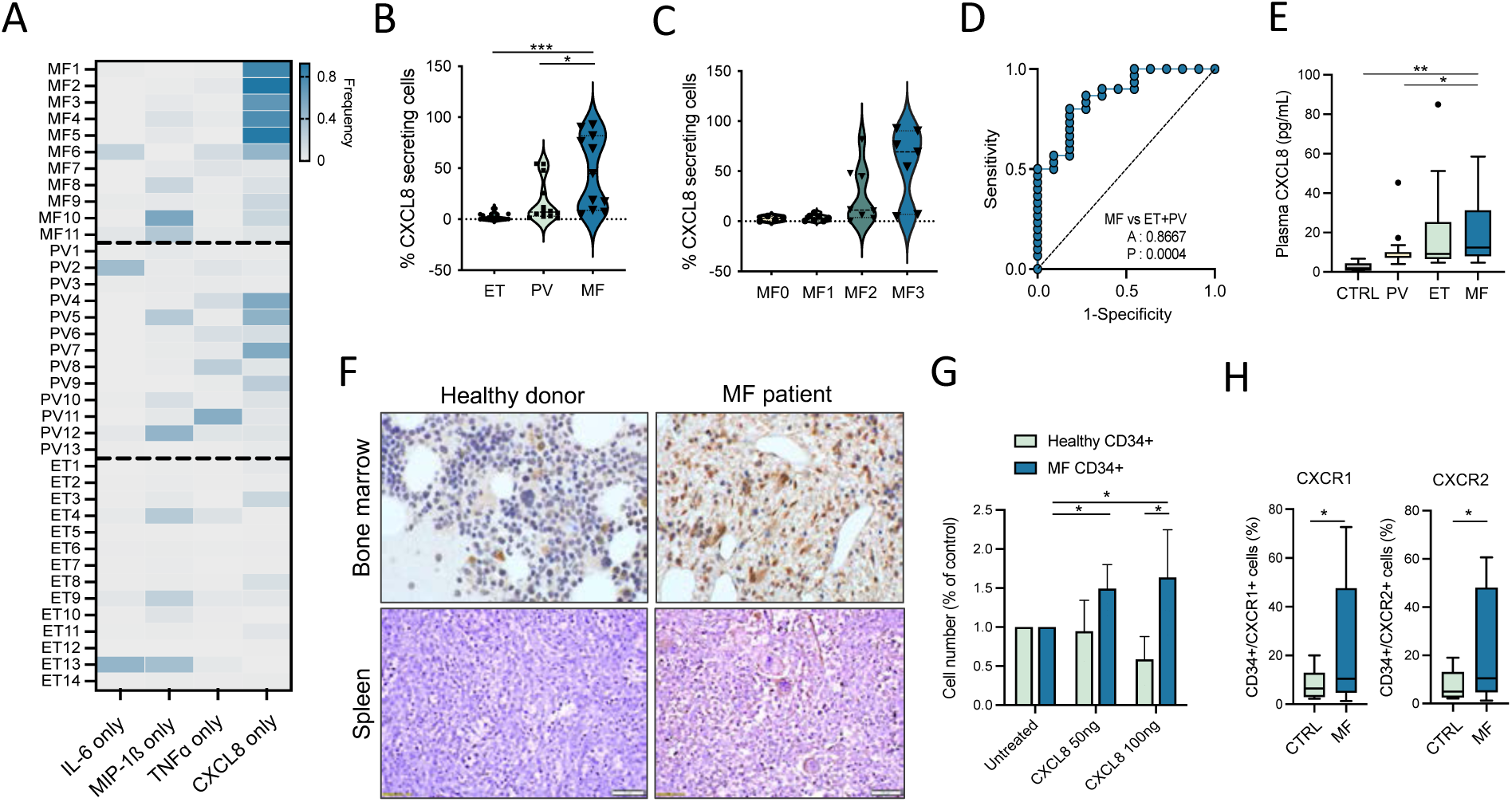
CXCL8-only secreting CD34+ cells are enriched in a subset of MF patients, and this correlates with clinical features, including grade reticulin fibrosis. **(A)** Heatmap demonstrating frequency of individual cytokine secreting CD34+ cells detected among individual patients across MPN sub-types myelofibrosis (MF), polycythemia vera (PV), and essential thrombocythemia (ET) as a percentage of total cytokine-secreting cells (0% in gray to 100% in dark blue). Four cytokines presented: IL-6, MIP-1β, TNFα, and CXCL8. **(B)** Violin plot depicting correlation between MPN sub-type and percent fraction of CXCL8-only secreting cells as detected by single-cell cytokine analysis. *p<0.05. ***p<0.001. **(C)** Violin plot depicting correlation between bone marrow reticulin score (MF0-3) of MF patients and percent fraction of detected CXCL8-only secreting cells by single-cell cytokine analysis. **(D)** Receiver operator curve demonstrating association between percent CXCL8-only secreting cells and MF sub-type in comparison to PV/ET. A=area under the curve; P=p-value. **(E)** CXCL8 levels as detected by ELISA in isolated plasma from PV, ET, and MF MPN patients in comparison to healthy controls (CTRL). (CTRL N=8, PV N=16, ET N=16, MF N=35). *p<0.05. **p<0.01. Data shown represent mean ± SD. **(F)** Representative CXCL8 immunohistochemistry (IHC) of healthy donors and MF bone marrow (BM) and spleen (from N=0/4 healthy donors and N=8/15 MF patients). **(G)** Ratio of total cell output relative to untreated of cultured healthy donor (HD; light blue) vs. MF (dark blue) CD34+ cells in response to two exogenous CXCL8 doses (50ng, 100ng). *p<0.05. Representative of triplicate experiments from N=3 HD and N=6 MF samples. Data shown represent mean ± SD. **(H)** Percent of total CD34+ cells expressing CXCR1 (left panel) or CXCR2 (right panel) by flow cytometry of control (CTRL) vs. MF patients (N=13 CTRL, H=15 MF). *p<0.05. Data shown represent mean ± SD.

We next investigated if CXCL8/CXCR2 potentiated signaling and enhanced fitness of MF HSPCs. MF CD34+ cells cultured in the presence of increasing doses of CXCL8 ligand *in vitro* showed dose-dependent proliferation (**Figure 2G**) as well as expansion of both CD33+ monocytic and CD41+ megakaryocytic cell numbers **(Supp. Figure 3C**). By contrast, WT CD34+ cells displayed reduced growth with increasing doses of CXCL8. Consistent with the sensitivity to CXCL8, we also detected increased surface expression of the CXCL8 receptors CXCR1 and CXCR2 on MF CD34+ cells by flow cytometry (**Figure 2H**), and CFU response to CXCL8 ligand correlated with degree of surface receptor expression **(Supp. Figure 3D)** suggestive of a positive feedback, autocrine effect. Of note, both CXCR1/2 surface expression and CXCL8 single-cell cytokine enumeration also correlated with JAK2^V617F^ variant allele frequency (VAF) **(Supp. Figure 3E-F)**, consistent with recent single cell studies,^36^ suggesting the degree of constitutive JAK/STAT activity corresponds with CXCL8/CXCR2 signaling output. Together with the above, these data suggested that CXCL8 is enriched and aberrantly secreted by multiple cell populations in MF and promotes cell growth and proliferation of MF HSPCs.

### Bulk integrated ATAC/RNA sequencing reveals enriched pathways in CXCL8 secretor vs. CXCL8 non-secretor MPN patients

We next wondered if transcriptional and chromatin states varied in the context of CXCL8 secretion. We performed bulk RNA sequencing (RNA-Seq) and Assay for Transposase-Accessible Chromatin with high-throughput sequencing (ATAC-Seq) on circulating CD34+ cells isolated from MPN patients with varying degrees of fibrosis and stratified the clinical samples into CXCL8 secretors vs. CXCL8 non-secretors based on their single-cell cytokine secretion profiles **(Supp. Table S3)**. UMAP visualization of RNA-Seq results revealed general clustering of CXCL8-secretors vs. non-secretors, which correlated generally with the degree of marrow fibrosis **(Supp. Figure 4A)**.

We identified 1,445 genes that were significantly upregulated and 529 genes that were significantly downregulated in CXCL8 secretor vs. CXCL8 non-secretor patient cells **(Supp. Figure 4B)**. Accessibility at the *CXCL8* locus correlated with the single-cell cytokine secretion data consistent with active *CXCL8* expression (**Figure 3A**). Review of the most differentially expressed genes (DEGs) by RNA-Seq alone between CXCL8 secretors vs. non-secretors revealed significant enrichment in genes encoding neutrophil markers and those involved in the activated innate immune response (e.g. *CTSG*, *ISG15, AZU1*, *MPO*, *PRTN3*, *ELANE*, and *RNASE2/3*) suggestive of a population skewed towards enhanced myeloid differentiation (**Figure 3B**). Consistent with our scRNA-Seq, gene set enrichment analysis (GSEA) confirmed significant enrichment in TNFα via NF-κB and Hallmark IFNα/*γ* response gene sets (**Figure 3C**), and network analysis revealed other gene ontology processes indicative of mature myeloid/neutrophil differentiation/activation and Toll-like Receptor (TLR) signaling (**Figure 3D**), increasingly implicated in MF.^11,21,37^ Integrated analysis of gene expression and chromatin accessibility data in a subset of patients for whom combined RNA- and ATAC- analysis was possible (N=5; **Supp. Table S3)** confirmed increased expression/accessibility of genes involved in the acute innate immune response, neutrophil activation/differentiation, and TLR signaling (e.g. the alarmins *S100a8/a9*, *CCL3*, *HCK*, *KLF4*, *CEBPB*, *TLR4*, *TRPM2*), as well as FOS/JUN activation (e.g. *FOS/FOSB*), extracellular matrix (ECM) remodeling (e.g. *PLAUR*), and type I IFNα response (including *IRF1/2/3*, *ELF3*, *OAS1/L*, *IFITM1/2*, *IFI6*, *LY6E*, and *IFI27*) (**Figure 3E**). These observations, coupled with increased expression of known reciprocal negative regulators of these pro-inflammatory pathways, including *DUSP1/2*, *ZFP36*, and *SOCS3*, suggest a MF CD34+ population skewed to neutrophil/monocytic differentiation, alarmin over-expression, TLR signaling, and neutrophil/monocyte activation. Notably, FOS/JUN (AP-1), like NF-κB, is a key regulatory factor that binds the *CXCL8* promoter to potently induce transcription^31^ suggesting combined signals at the *CXCL8* promoter act to modulate its activation.

**Figure 3:**
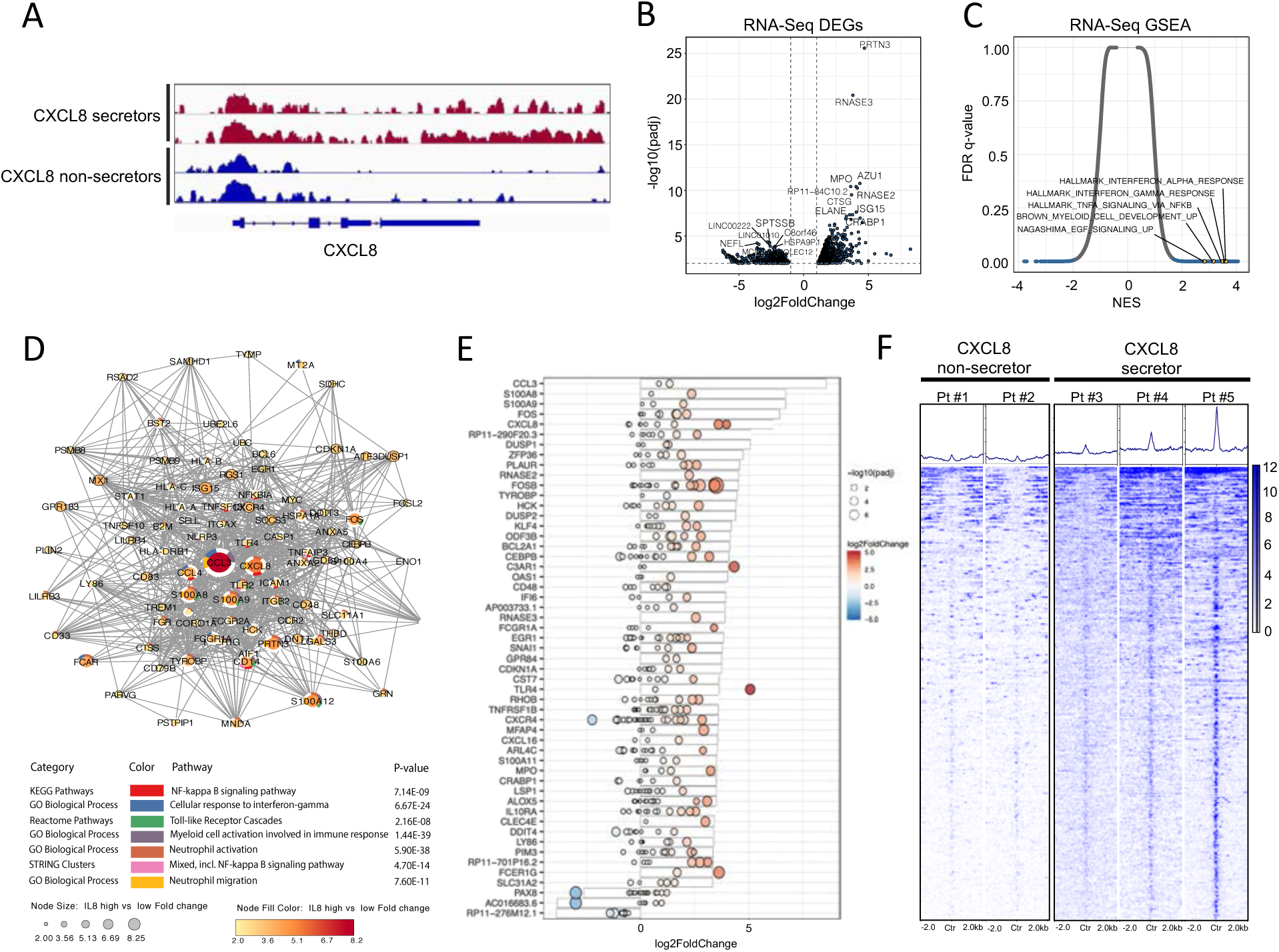
Integrated transcriptional (RNA-Seq)/chromatin accessibility (ATAC-Seq) profiling identifies pathways enriched in CXCL8-secretor MF. **(A)** Representative accessibility peaks at the *CXCL8* locus of representative CXCL8-secretor vs. non-secretor MPN patients. **(B)** Volcano plot demonstrating most differentially expressed genes (DEGs) in CXCL8 secretor vs. non-secretor MPN patients by RNA-Seq. The significant events with an inclusion level >0.5 log fold change and an FDR-corrected P<0.0001 are shown in blue. **(C)** Gene Set Enrichment Analysis (GSEA) demonstrating enriched pathways of CXCL8 secretors vs. non-secretors by RNA-Seq plotted as normalized enrichment score (NES) by FDR q-value. **(D)** Optimized gene expression sub-network analysis from gene expression profiles showing key nodes enriched by RNA-Seq in CXCL8 secretor vs. non-secretor MPN patients. Additional information regarding the creation of sub-networks can be found in the STAR Methods. **(E)** Waterfall plot with integrated gene expression and chromatin accessibility showing most differentially regulated genes (represented as log2FoldChange) in CXCL8-secretors vs. non-secretors and their corresponding degree of changes in accessibility peaks (represented as log2FoldChange and -log10(padj), red, positive values; blue, negative values). **(F)** Tornado plot and heatmaps depicting accessibility at promoter regions of the top 500 leading edge genes in the TNFα via NF-κB gene set of CXCL8 non-secretor (N=2) vs. CXCL8 secretor (N=3) MPN patients (see Supp. Table S3).

Given the degree of enrichment in TNFα via NF-κB in CXCL8-secretor patients and previous associations of this gene set/pathway in MF, we sought to determine if the chromatin state of CXCL8-secreting MF represented a distinct pro-inflammatory MPN entity or whether all MPN CD34+ cells were poised for inflammatory signaling through NF-κB irrespective of their CXCL8 secretion level. Assessment of the accessibility landscape surrounding the leading edge genes which were most responsible for driving the Hallmark TNFα and IFNα/*γ* pathways revealed a marked increase in ATAC signal in CXCL8-secretor compared with CXCL8-non-secretor patient samples, concordant with their transcriptional output (**Figure 3F**). This suggests that, irrespective of MPN sub-type, rather than being inherently poised for NF-κB inflammatory signaling, additional epigenetic changes are required to engage this pro-inflammatory program.

To validate our internal gene expression/accessibility data and uncover additional gene pathways enriched in CXCL8 secretors vs. non-secretors, we extended our analysis to a larger (N=39) publicly available transcription microarray dataset of MF CD34+ cells^38^ which we stratified by *CXCL8* expression level into upper 20% and lower 20% thresholds for differential expression analysis **(Supp. Figure 4C)**. Notably, we observed strong concordance with our internal dataset **(Supp. Figure 4D)**, and hierarchical clustering analysis revealed similar upregulation of key pro-inflammatory TLR receptor genes, including neutrophil markers (*FCN1*, *LYZ*), the alarmins *S100a8/a9*, TLR signaling elements (*LY96*, *TLR8*) as well as pro-inflammatory transcription factors *FOS* and *JUN* **(Supp. Figure 4E)**. GSEA analysis of the external cohort revealed similar enrichment in TNFα via NF-κB and IFNα/*γ* pro-inflammatory gene sets as well as other gene sets associated with mature myeloid cell activation and TLR signaling **(Supp. Figure 4F-G)**. These data, together with our internal cohort, establish an important role for neutrophil activation and acute phase inflammatory response in CXCL8-secreting MF CD34+ cells and highlight how stratification of expression patterns by CXCL8 alone might underscore this phenotype.

### *Cxcr2* deletion improves hematologic parameters and reduces fibrosis in the hMPL^W515L^ adoptive transfer model of MF

We next sought to functionally validate the role of the CXCR2 pathway in fibrosis development using the hMPL^W515L^ bone marrow fibrosis mouse model.^39^ While mice lack the functional analog equivalent of human CXCL8,^40^ the murine CXCR1/2 receptors are highly homologous to that of human and have been shown to bind human CXCL8 and activate similar downstream mediators.^41^ Thus, we reasoned this was an appropriate *in vivo* system to assess CXCR2 pathway signaling in the context of mutant-driven fibrosis. Consistent with this, culture of cKit+ murine bone marrow cells in the presence of recombinant human CXCL8 demonstrated enhanced signaling through pERK and pSTAT3 that was abrogated in the setting of *Cxcr2* knock-out (KO) **(Supp Figure 5A)**.

We thus investigated the effects of genetic deletion of *Cxcr2* within the murine bone marrow hematopoietic compartment on MPL^W515L^-driven fibrosis development. Isolated *VavCre-Cxcr2^-/-^* or Cre-negative (wild-type; WT) *Cxcr2^f/f^* bone marrow cells were transduced with MSCV-hMPL^W515L^-IRES-GFP and transplanted into lethally-irradiated recipient mice and monitored for the development of MF **(Supp. Figure 5B)**. *Cxcr2* knock-out was validated by loss of surface expression on mature myeloid cells in primary *VavCre-Cxcr2^-/-^* mice **(Supp. Figure 5C)** and in transplant recipients. Mice transplanted with *Cxcr2^-/-^* hMPL^W515L^-expressing cells displayed significant improvements in white blood cell (WBC; mean 163 K/uL vs. 23 K/uL, p<0.01) and platelet parameters (mean 596 K/uL vs. 113 K/uL, p<0.05) compared to Cre-negative (WT) *Cxcr2^f/f^*-expressing hMPL^W515L^ mice (**Figure 4A**). Further, CD11b+Gr1+ neutrophil fractions were also significantly reduced in the peripheral blood of *Cxcr2^-/-^* hMPL^W515L^ mice **(Supp.Figure 5D-E)**, this in contrast to the enhanced neutrophilia observed with primary *Cxcr2^-/-^* knock-out mice^42,43^ suggesting a paradoxical/reciprocal effect on neutrophils in the setting of hMPL^W515L^. In addition to improvements in blood cell count parameters, mutant GFP+ cell fraction (mean 75% vs. 12.5%, p<0.0001) (**Figure 4B**) as well as total spleen weights (mean 538mg vs. 295mg, p<0.01) (**Figure 4C**) were also significantly reduced in *Cxcr2^-/-^* hMPL^W515L^ mice, with concomitant reductions in extent of extramedullary hematopoiesis (EMH). Most importantly, histopathologic analysis revealed significant reductions in the degree of reticulin fibrosis in both the bone marrow and spleen **(Figure 4D, Supp. Figure 5F-G)**, which, together with the findings above, was associated with a significant improvement in overall survival of *Cxcr2^-/-^* hMPL^W515L^ mice in comparison to *Cxcr2^f/f^* WT hMPL^W515L^ mice (median 84d vs. 42d, p<0.01) (**Figure 4E**).

**Figure 4:**
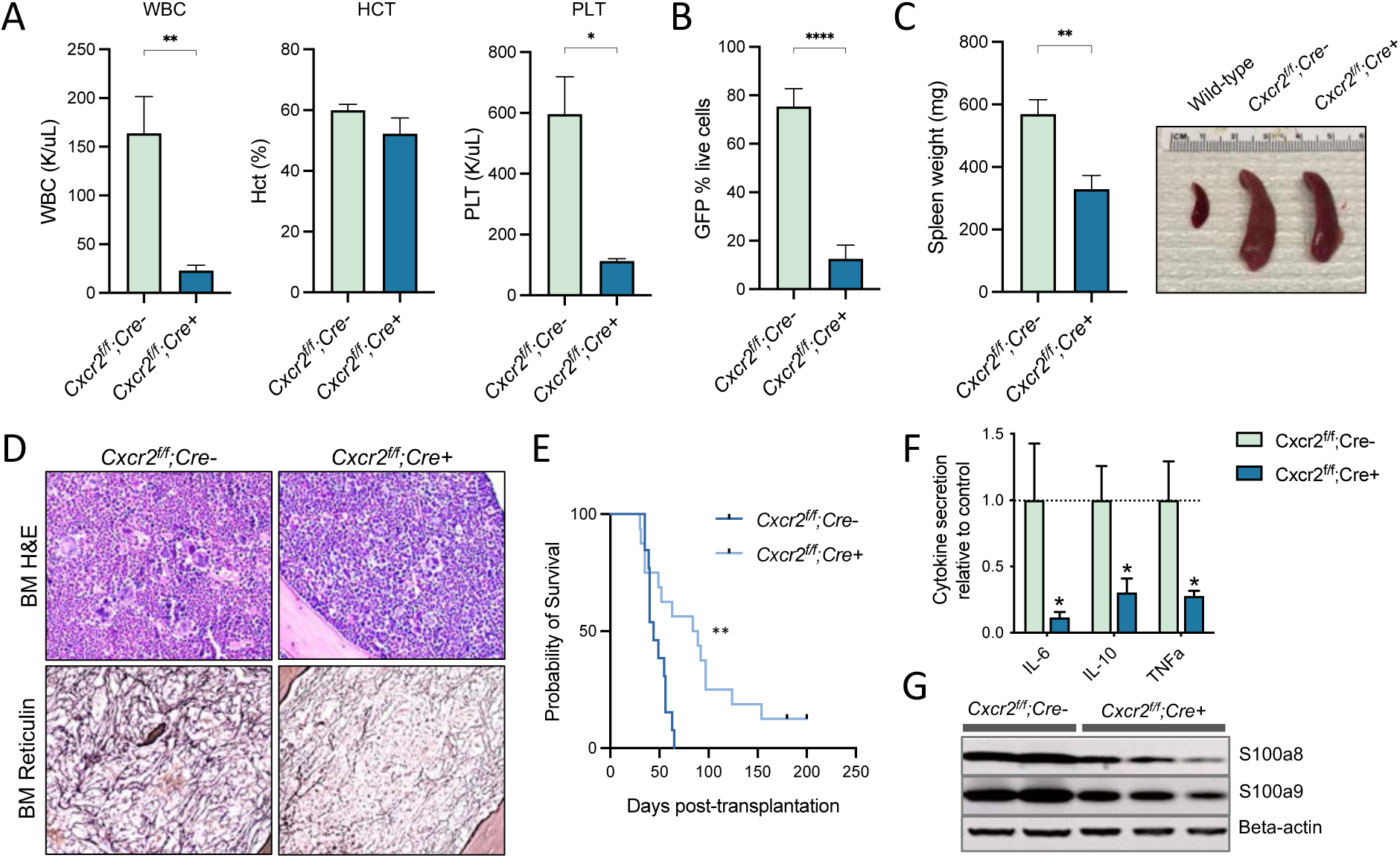
*Cxcr2* deletion in murine bone marrow improves counts and reticulin fibrosis in the hMPL^W515L^ adoptive transfer model of myelofibrosis. **(A)** White blood cell counts (WBC, K/uL), hematocrit levels (Hct, %), and platelet counts (PLT, K/uL) of *Cxcr2^f/f^;Cre^+^* knock-out (KO) hMPL^W515L^ mice compared to *Cxcr2^f/f^;Cre^-^* wild-type (WT) hMPL^W515L^ mice. *p-value <0.05. **p-value <0.01. Data shown represent mean ± SEM. The Student’s test (unpaired, two-tailed) was used to compare the mean of two groups. **(B)** Peripheral blood mutant cell fraction by green fluorescent reporter (GFP) percentage in *Cxcr2^f/f^;Cre^+^* hMPL^W515L^ mice vs. *Cxcr2^f/f^;Cre^-^* wild-type (WT) hMPL^W515L^ mice. ****p-value <0.0001. Data shown represent mean ± SEM. **(C)** Spleen weights (mg) and representative spleens of *Cxcr2^f/f^;Cre^+^* KO vs. *Cxcr2^f/f^;Cre^-^* WT hMPL^W515L^ mice. **p-value <0.01. Data shown represent mean ± SEM. **(D)** Representative hematoxylin and eosin (H&E) and reticulin images of bone marrow from *Cxcr2^f/f^;Cre^+^* vs. *Cxcr2^f/f^;Cre^-^* WT hMPL^W515L^ mice. 20X magnification. **(E)** Kaplan-Meier survival analysis of *Cxcr2^f/f^;Cre^+^* KO hMPL^W515L^ mice vs. *Cxcr2^f/f^;Cre^-^* WT hMPL^W515L^ mice. **p-value <0.01 (log-rank test). **(F)** Fold change in serum cytokine levels of IL-6, IL-10, and TNFα of *Cxcr2^f/f^;Cre^+^* KO compared with *Cxcr2^f/f^;Cre^-^* WT hMPL^W515L^ mice. N=8/arm. *p<0.05. Data shown represent mean ± SEM. **(G)** Western blot analysis of the alarmins S100a8/a9 from harvested splenocytes of *Cxcr2^f/f^;Cre^+^* KO vs. *Cxcr2^f/f^;Cre^-^* WT hMPL^W515L^ mice.

Megakaryocytic hyperplasia and proliferation are pathognomonic features of myelofibrosis in MPN and are thought to promote elicitation of various pro-fibrogenic cytokines, including CXCL4 and TGFβ.^29,44^ Our *in vitro* MF primary mononuclear cell culture data suggested that CXCL8 can directly influence megakaryocytic output/proliferation. Given this, we evaluated the effect of *Cxcr2* loss on CD41+ megakaryocyte fractions and degree of megakaryocytic hyperplasia in the hMPL^W515L^ model. Flow cytometric analysis revealed significant reductions in the percentage of CD41+ cells in bone marrow (mean 7.5% vs. 1%, p<0.01) **(Supp. Figure 5H)**, and histopathologic sections of bone marrow of *Cxcr2^-/-^* hMPL^W515L^ recipient mice confirmed reductions in observable megakaryocytes **(Supp. Figure 5I)**.

Finally, to assess the effect of *Cxcr2* loss on pro-inflammatory cytokine enumeration and TLR signaling, we assessed secreted pro-inflammatory cytokine levels in serum of transplanted hMPL^W515L^ mice by Luminex bead-based cytokine profiling. Notably, *Cxcr2^-/-^* hMPL^W515L^ mice displayed significant reductions in critical TLR-mediated cytokines, specifically IL-6, IL-10, and TNFα in comparison to *Cxcr2^f/f^* WT hMPL^W515L^ mice (**Figure 4F**). In further support of this, western blot analysis of harvested splenocytes from *Cxcr2^-/-^* hMPL^W515L^ mice revealed reductions in detectable levels of the TLR agonists S100a8/a9 (**Figure 4G**). These data are consistent with our *in vitro* and RNA-Seq transcriptional patient data and further suggest a role for CXCR2 in modulating TLR-mediated pro-inflammatory pathway signaling.

### Pharmacologic inhibition of CXCR1/2 improves peripheral blood counts and reduces fibrosis in the hMPL^W515L^ model of myelofibrosis

We next sought to validate our genetic deletion studies and assess CXCR1/2 signaling as a potential therapeutic target in MF by evaluating pharmacologic inhibition of the CXCR1/2 pathway in the hMPL^W515L^ model. After hMPL^W515L^ transplanted mice displayed evidence of disease, including leukocytosis, inflammatory cytokine production, and BM fibrosis, mice were assigned based on WBC count into four separate treatment arms: vehicle, ruxolitinib (60mg/kg orally BID), the CXCR1/2 inhibitor reparixin (60mg/kg SubQ BID), or combination therapy **(Supp. Figure 6A)**. No significant weight loss was observed across reparixin treatment arms during the 3-week treatment period compared to vehicle treated mice **(Supp. Figure 6B)**. Consistent with our genetic knock out studies, mice treated with reparixin, either alone or in combination with ruxolitinib, demonstrated improvements in leukocytosis (mean 251 K/uL vehicle vs. 80.4 K/uL reparixin vs. 30.7 K/uL combo, p<0.05) and platelet counts (mean 3437.8 K/ul vehicle vs. 1244.2 K/uL reparixin vs. 742 K/uL combo, p<0.05) in comparison to vehicle treated mice (**Figure 5A**). Reparixin alone however led to more modest, non-significant reductions in spleen weights (mean 585.8mg vs. 490.5mg) **(Figure B)**, and, like ruxolitinib, had only minimal effects on percent Mac1+Gr1+ peripheral blood neutrophil (mean 75.2% vs 67.9%, p=0.3) fraction **(Supp. Figure 6C)**. Of note, significant reductions in megakaryocytic number were observed in bone marrow with reparixin treatment, consistent with our genetic knock-out studies (**Figure 5C**). Most importantly, reparixin-treated mice exhibited significant reductions in the degree of reticulin fibrosis, both in the bone marrow and spleen **(Figure 5D-E; Supp. Figure 6D)** that was further enhanced by combination treatment with ruxolitinib. These data suggest targeted inhibition of the CXCR1/2 pathway alone is sufficient to ameliorate fibrosis in a JAK/STAT driven model of fibrosis and is synergistic with JAK inhibitor therapy.

**Figure 5:**
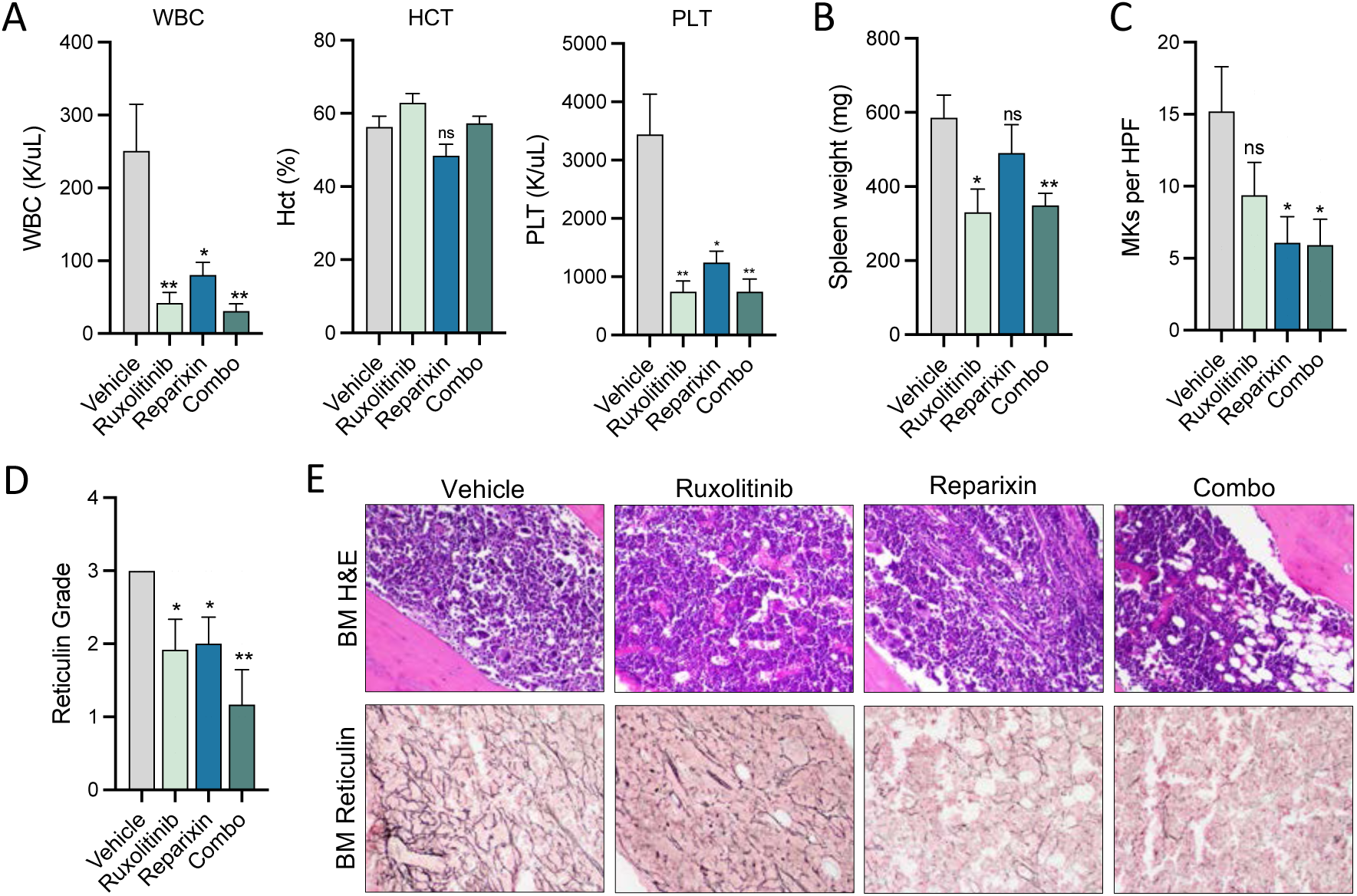
Pharmacologic inhibition of CXCR1/2 improves hematologic parameters and reticulin fibrosis in the hMPL^W515L^ adoptive transfer model of myelofibrosis. **(A)** White blood cell counts (WBC, K/uL), hematocrit levels (Hct, %), and platelet counts (PLT, K/uL) of hMPL^W515L^- diseased mice treated with vehicle, ruxolitinib (60mg/kg twice daily), the CXCR1/2 inhibitor reparixin (60mg/kg twice daily), or combination therapy at timed sac following 21 days of treatment. *p-value <0.05. **p-value <0.01. ns=not significant. The Student’s test (unpaired, two-tailed) was used to compare the mean of two groups. Data shown represent mean ± SEM. N=6/condition. **(B)** Spleen weights (mg) of hMPL^W515L^-diseased mice treated with either vehicle, ruxolitinib, reparixin, or combination therapy. *p-value <0.05. **p-value <0.01. ns=not significant. Data shown represent mean ± SEM. **(C)** Megakaryocyte number (MKs) per high powered field (HPF) observed in hMPL^W515L^ mice in response to treatment. *p-value <0.05. ns=not significant. Data shown represent mean ± SEM. **(D)** Bone marrow reticulin scores of hMPL^W515L^-diseased mice treated with either vehicle, ruxolitinib, reparixin, or combination therapy. *p-value <0.05. **p-value <0.01. N=6/condition. **(E)** Representative H&E and reticulin images of hMPL^W515L^-diseased bone marrow treated with ruxolitinib, reparixin, or combination therapy compared with vehicle-treated mice. 20X magnification.

### CXCR1/2 inhibition ameliorates bone marrow fibrosis in the *Gata1^low^* model of MF

We next evaluated therapeutic efficacy of CXCR1/2 inhibition with reparixin in the *Gata1^low^* model of MF.^45^ Because *Gata1^low^* mice develop MF independent of constitutive JAK/STAT activity, we reasoned this model could be used to validate the role of the CXCR2 pathway in bone marrow fibrosis development irrespective of JAK/STAT mediated inflammatory signaling. Increased expression of Cxcl1, as well as the Cxcr1/2 receptors was validated in *Gata1^low^* mice as was observed in MF patients **(Figure 6A, Supp. Figure 7A)**. 8-month old mice with emerging fibrotic features were then treated with vehicle (sterile saline SubQ) or reparixin (7.5mh/h/Kg SubQ by continuous infusion) for 3 weeks and then evaluated for treatment response. Once again, no significant weight changes were seen throughout the treatment period **(Supp. Figure 7B)**. Changes in blood counts in response to reparixin therapy were more modest in *Gata1^low^* mice (**Figure 6B**), perhaps consistent with the generally milder degree of disordered blood counts often observed with these mice in comparison to the hMPL^W515L^ model; nor were there any significant changes in spleen volumes or bone marrow cellularity in response to treatment **(Supp. Figure 7C-D)**. Strikingly however, reparixin-treated *Gata1^low^* mice displayed a dramatic reduction in the level of reticulin fibrosis in both bone marrow and spleen sections **(Figure 6C, Supp. Figure 7E-F)**, and were confirmed when the reticulin fibers were independently evaluated by Gomori staining **(Supp. Figure 7G)**. Together, these data confirm an important role for CXCR2 pathway signaling in bone marrow fibrosis progression and validate CXCR1/2 as a potential target for the treatment of bone marrow fibrosis irrespective of MPN driver mutation status.

**Figure 6:**
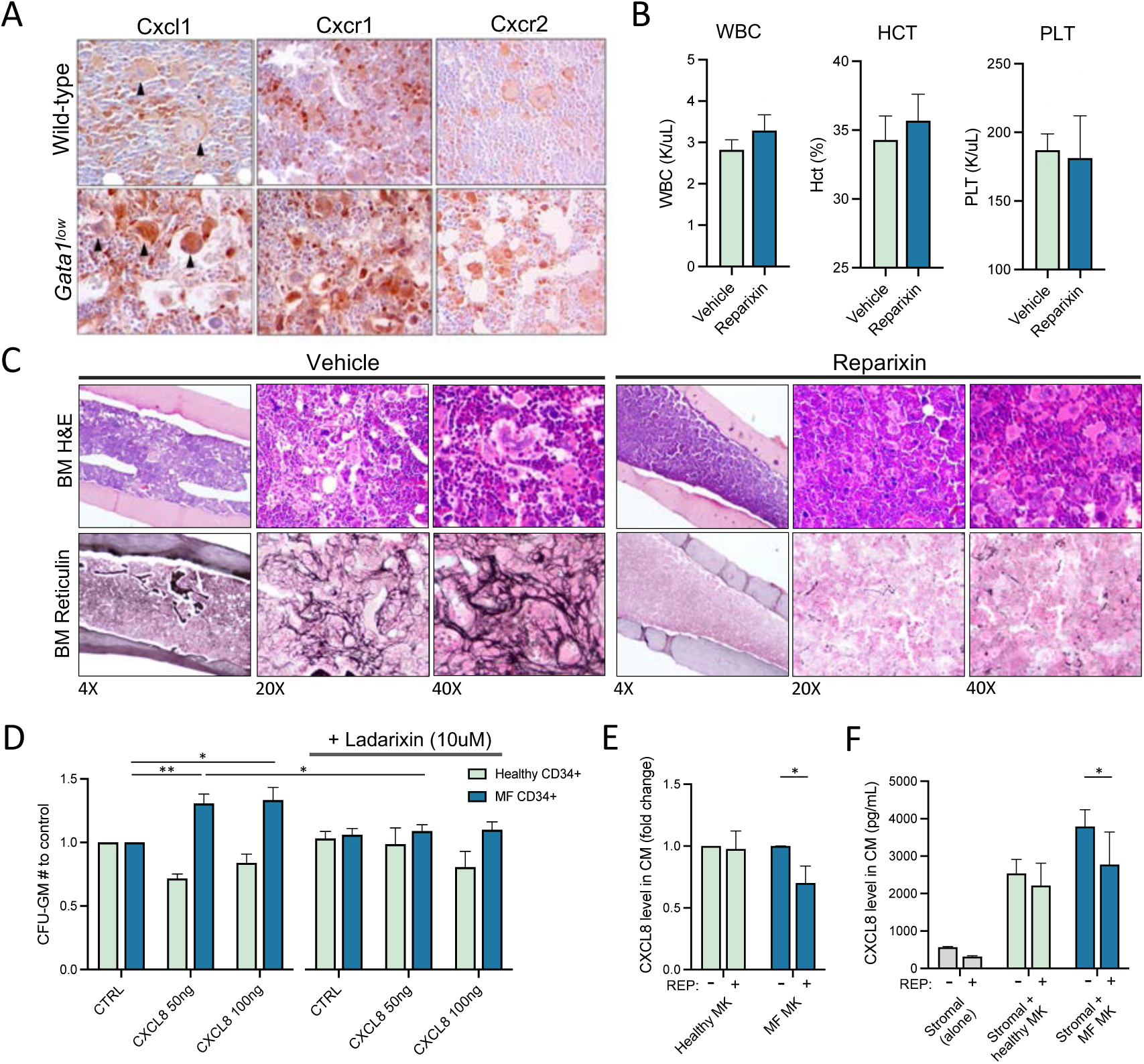
CXCR1/2 inhibition demonstrates therapeutic efficacy in the *Gata1^low^* model of myelofibrosis. **(A)** Representative Cxcl1, Cxcr1 and Cxcr2 immunohistochemistry of *Gata1^low^* mice with active fibrosis in comparison to age-matched wild-type controls. N=6/arm. 40X magnification (quantitative data are presented in Supp. Figure 7A). **(B)** White blood cell counts (WBC, K/uL), hematocrit levels (Hct, %), and platelet counts (PLT, K/uL) of *Gata1^low^* mice treated with reparixin compared to mice treated with vehicle. N=3-4/condition. Data shown represent mean ± SEM. **(C)** Representative H&E and reticulin images of *Gata1^low^* bone marrow treated with reparixin therapy compared to vehicle-treated mice. Magnification as indicated. **(D)** Colony forming unit (CFU) assay demonstrating total granulocyte-macrophage progenitor (CFU-GM) colony number relative to control of cultured healthy donor (HD; light blue) vs. MF (dark blue) CD34+ cells with exogenous CXCL8 ligand and response to the second-generation CXCR1/2 antagonist Ladarixin (10uM) *in vitro*. *p<0.05. **p<0.01. Representative of duplicate experiments from 5 HD and 13 individual MF cases. **(E)** Fold-change in detectable CXCL8 levels in conditioned media (CM) elicited by either healthy donor vs. MF megakaryocytes (MKs) with or without the addition of reparixin (REP; 10uM). *p-value <0.05. Data shown represent mean ± SD. **(F)** Total levels of CXCL8 in conditioned media (CM) of cultured stromal cells, either alone or together with healthy vs. MF megakaryocytes (MKs) with or without the addition of reparixin (REP; 10uM). *p-value <0.05. Data shown represent mean ± SD.

### CXCR1/2 inhibition demonstrates efficacy against primary MPN cells *in vitro*

We next assessed the impact of CXCR1/2 inhibitor therapy on the proliferation and colony forming capacity of primary MF CD34+ progenitor cells *in vitro*. Consistent with our liquid culture experiments above, CD34+ cells from MF patients demonstrated enhanced colony forming capacity in the presence of increasing doses of exogenous CXCL8 highlighting the role of CXCL8 in HSC clonal outgrowth. Notably, this effect was abolished with the addition of ladarixin, a second generation CXCR1/2 inhibitor (**Figure 6D**). Similar effects were also observed with CD33+ and CD41+ cell output in response to therapy **(Supp. Figure 8A-B)** consistent with our *in vivo* studies. Furthermore, treatment with reparixin also reduced levels of both CXCL8 and VEGF elaborated by cultured MF megakaryocytes **(Figure 6E, Supp. Figure 8C)** suggesting downregulation of an autocrine feedback loop.

To further evaluate the effects of CXCL8 on MF malignant clones, we performed *JAK2^V617F^* genotyping using nested PCR of isolated MF hematopoietic colonies cultured with or without CXCL8. Specifically, individual colonies were plucked from six different cases of *JAK2^V617F^*^+^ MF and then genotyped. Treatment with CXCL8 increased the absolute number of *JAK2^V617F+^* colonies in all six cases, including in 2 cases with undetectable *JAK2^V617F+^* colonies compared to the control group **(Table 1)**. The effects of enhanced *JAK2^V617F+^* mutant colony output in the setting of exogenous CXCL8 however were blocked by administration of Ladarixin in 4/6 MF cases.

**Table 1.**
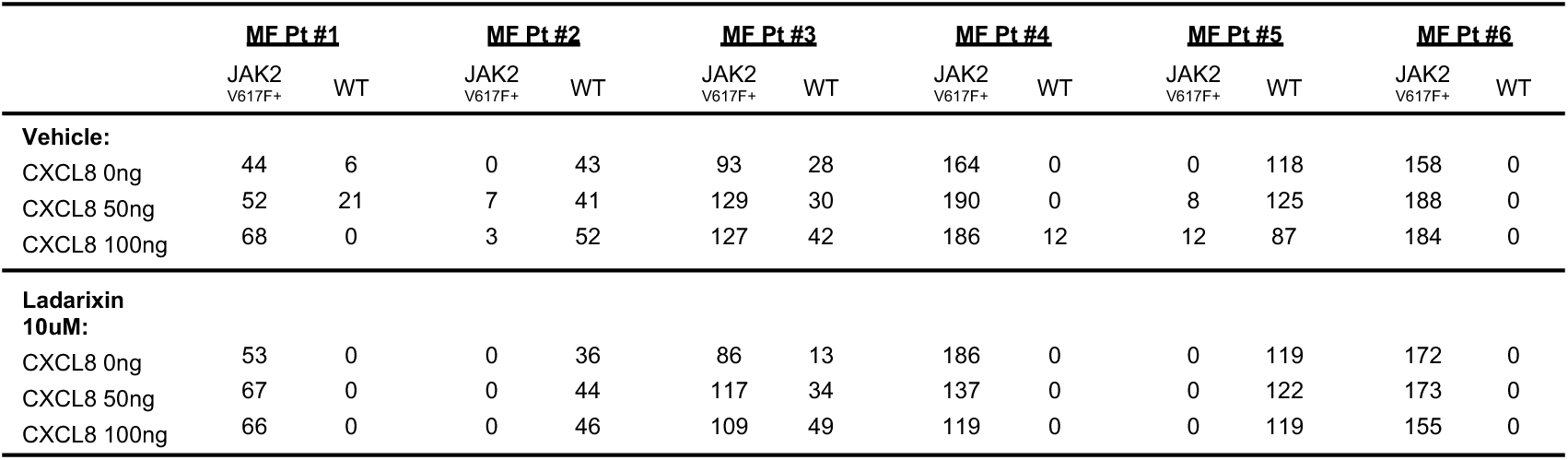
JAK2^V617F^ colony genotyping data in response to exogenous CXCL8 +/- CXCR1/2 inhibition.

Finally, we evaluated CXCL8 enumeration by MF megakaryocytes in the presence of bone marrow stromal cells using a previously described co-culture method.^46^ Notably, a greater abundance of secreted CXCL8 and VEGF was observed when stromal cells and isolated MKs were co-cultured together than when cultured separately, and MF MKs, when co-cultured with stromal cells, produced an even greater degree of CXCL8 and VEGF than WT MKs **(Supp. Figure 8D-E)**. Notably, this effect too was abrogated when cells were treated with reparixin **(Figure 6F, Supp. Figure 8F)**. Together, these data suggest that the elaboration of CXCL8 is the consequence of autocrine and paracrine regulatory loops involving HSCs, MKs, neutrophils, and stromal cells in MF, and that these loops can be interrupted by CXCR1/2 inhibition.

## DISCUSSION

Aberrant pro-inflammatory signaling is a hallmark feature of MPNs.^19^ Therapy with current JAK1/2 inhibitors improves symptoms and clinical outcomes in MF underscoring the role of constitutive JAK/STAT and pro-inflammatory signaling in disease maintenance.^47–49^ While many pro-inflammatory cytokine levels are reduced with JAK inhibitor therapy, others, including CXCL8, are not,^37,47^ suggesting that alternative sustained pro-inflammatory pathways play a critical role in myelofibrotic progression. Previously we and others identified an NF-κB pro-inflammatory signaling network that promotes MF.^24,25,37^ Additional understanding of how specific cytokine mediators drive fibrosis progression through canonical NF-κB will provide important insights into mechanisms contributing to fibrosis development and identify biomarkers predictive of treatment response. Given emerging data highlighting the role of individual cytokine signaling in MF progression,^29,37^ we sought to determine specific cytokine pathways implicated through NF-κB that contribute to this process.

In this study, we determined that the CXCL8-CXCR2 axis is an important mediator of fibrosis progression by utilizing individual primary MF patient samples and two separate murine models of MF. CXCL8 is one of many pro-inflammatory cytokines up-regulated by NF-κB signaling and is a potent inducer of neutrophil differentiation and mobilization to sites of acute infection. Aberrant CXCL8-CXCR2 signaling is implicated in numerous pro-inflammatory conditions and autoimmune phenomena, including solid organ fibrosis.^50,51^ In epithelial malignancies, CXCL8 is thought to enhance neoangiogenesis and extracellular matrix (ECM) remodeling and consequently promote a tumor microenvironment conducive for furthered growth and metastatic progression.^52^ In hematologic malignancies, CXCL8 promotes leukemic stem cell (LSC) fitness in chronic myelogenous leukemia (CML), myelodysplastic syndrome (MDS), and acute myeloid leukemia (AML).^53,54^ Notably, CXCR1/2 antagonists induce apoptosis of AML LSCs *in vitro.*^53,55^ In the present study, we used integrated single-cell transcriptional and cytokine assays across multiple MPN sub-types and validate strong enrichment of a CXCL8 signature in MF in comparison to pre-fibrotic MPN sub-types PV and ET. Notably, our transcriptional data suggests that CXCL8-CXCR2 signaling is enhanced across the hematopoietic hierarchy in MF and aberrantly activated throughout myeloid lineage commitment. Furthermore, we demonstrate that MF CD34+ cells display dose-dependent cell growth in response to CXCL8. These data together suggest that MF cells are primed for, and elicit, on-going pro-inflammatory signaling through CXCR2. Importantly, our data suggest that the presence of CXCL8 secreting HSPCs might represent a circulating biomarker for bone marrow fibrosis, and prospective studies can delineate if emergence of a CXCL8-secreting clone over time predicts for fibrotic progression in MPN.

The myeloid differentiation program enriched in CXCL8-high MF is reminiscent of an acute hematopoietic stress response and adds to expanding literature on the role of Toll like receptor (TLR) signaling in MF and myeloid disease.^37,56,57^ We previously have shown that NF-κB is activated in both mutant and non-mutant hematopoietic cells in MPL^W515L^-diseased mice *in vivo*,^35^ and other studies have demonstrated that HSCs express TLR receptors that potently up-regulate pro-inflammatory cytokines in response to various pathogen- and damage-associated molecular patterns (PAMPs/DAMPs), including the TLR agonists S100a8/a9, to promote myeloid cell maturation and mobilization in an autocrine and paracrine manner.^58,59^ In MF, HSPCs are preferentially sensitive to TLR agonists *in vitro*,^37^ and S100a8/a9 play key roles in mesenchymal stromal cell proliferation/differentiation to myofibroblasts and collagen deposition^11^ and have been implicated in fibrosis previously^60^ suggesting simultaneous roles for TLR signaling on MF HSPCs and their surrounding microenvironment. As stated, CXCL8 is a potent inducer of myeloid cell chemotaxis and activation. Our data suggest then that CXCL8 elicited by mutant HSCs and other myeloid cells promotes an early feed-forward loop of enhanced S100a8/a9 release and TLR signaling that over time, reinforces MSC transcriptional changes that favor fibroblastic proliferation. Importantly, our *in vivo* genetic deletion and pharmacologic studies suggest that this cycle can be disrupted by direct therapeutic inhibition of the CXCL8-CXCR2 axis with concomitant improvements in megakaryocyte and neutrophil numbers, extramedullary hematopoiesis, and fibrosis within the marrow and spleen.

In addition to NF-κB, we also previously identified enrichment in a HIF1α signature in MF mice,^61^ and enhanced HIF1α stabilization through loss of the *Lkb1* tumor suppressor was recently shown to enhance fibrosis development in the hMPL^W515L^ mouse model.^62^ Notably, CXCL8 itself promotes MSC migration under hypoxic conditions,^63^ and hypoxia was shown to induce expression of the lncRNA MALAT1, which promotes *CXCL8* expression directly by epigenetic alteration of the *CXCL8* locus by p300.^33^ These data together suggest that the CXCR2 signaling axis might serve as a key node through which multiple pro-inflammatory cytokines and hypoxia-mediated pathways act to promote fibrosis in MPNs. How these various pathways evolve in relation to constitutive JAK/STAT signaling then remains unclear. TPO signaling has been shown to induce production and release of Cxcr2 ligands Cxcl1 and Cxcl2 in mice to promote neutrophil mobilization/activation from the marrow.^64^ Further, STAT3 has been shown to modulate the degree of CXCR2 transcription and signaling by influencing ERK signal intensity.^65^ This might explain why JAK2^V617F^ homozygous MF cells demonstrate increased *CXCL8* expression and suggests perhaps these cells might be preferentially sensitive to CXCR1/2 inhibitor therapy. Still, that we observed enrichment of CXCL8-CXCR2 signaling in only a subset of MF patients suggests that likely additional factors, inflammatory signals, and adaptive changes over time converge to promote fibrosis in MPN.

In sum, our findings show that CXCL8, potently induced by NF-κB signaling, is a key mediator of fibrosis development in MF and represents an emerging important biomarker for fibrotic progression in MPNs. Our findings further support the role of activated innate immune signaling pathways in MF and add to existing literature establishing CXCL8-CXCR2 pro-inflammatory signaling as a critical node driving disease progression across the spectrum of myeloid disorders, including in MPN as well as in MDS and AML. Moreover, our studies suggest that inhibition of CXCL8/CXCR2 signaling represents a potential therapeutic opportunity to intercept MPN progression and bone marrow fibrosis, which should be further studied in the clinical context.

## AUTHOR CONTRIBUTIONS

A.D., D.K., M.L, A.R.M, R.F., R.L., and R.H. conceived the project, designed the experiments, and analyzed the data. ScRNA-Seq and single-cell cytokine sample processing and analysis were performed primarily by D.K., Z.C., Y.X, and R.F. *In vitro* work was performed primarily by M.L. with technical assistance from L.X. and N.E. hMPL^W515L^ and genetic knock-out mouse studies as well as patient sample collection/processing/clinical annotation were performed primarily by A.D. and M.F. with technical assistance from Y.P., A.K., J.S., A.K., and J.Y. *Gata1^low^* studies were performed primarily by F.G., P.V., and F.M. Bulk ATAC-Seq and RNA-Seq computational analyses were performed primarily by J.L.Y. and R.K.. R.R., E.M., M.K., M.S., J.C., E.T., J.Z., C.H., A.Z., K.C., and T.M assisted with management of clinical data and specimens. W.X assisted with pathological assessment of specimens. A.D., R.L.L, and R.H. contributed to the first draft of the manuscript. Funding acquisition: R.L.L., R.H., R.F., A.R.M.

## ACKNOWLEDGEMENTS

The authors thank members of the Levine laboratory for helpful comments and discussion, including Robert Bowman for critical insights and assistance with data presentation. We also thank the members of the Hoffman laboratory and MPN-RC tissue bank as well as ISMMS tissue bank for providing MF patients’ samples. Studies supported by MSK core facilities were supported in part by MSKCC Support Grant/Core Grant P30 CA008748 and the Marie-Josée and Henry R. Kravis Center for Molecular Oncology. This work was supported by National Cancer Institute awards P01 CA108671 (R. H., R.L.L., A. R. M.) and AIRC IG23525 (A.R.M.). R.L.L. was supported by a Leukemia and Lymphoma Society Scholar award. A.J.D. is a William Raveis Charitable Fund Physician-Scientist of the Damon Runyon Cancer Research Foundation (PST-24-19). He also has received funding from the American Association of Cancer Research and the American Association of Clinical Oncology. R.L.L. is also supported by a Leukemia & Lymphoma Society Specialized Center of Research grant. T.M. and J.Z. work on this project was supported by NIH SBIR grant R44EB023777. The personnel of the Animal Facilities of Istituto Superiore di Sanità are acknowledged for assistance with the *Gata1^low^* experiments.

## DECLARATION OF INTERESTS

R.L.L. is on the supervisory board of Qiagen and is a scientific advisor to Imago, Mission Bio, Bakx, Zentalis, Ajax, Auron, Prelude, C4 Therapeutics and Isoplexis. He has received research support from Abbvie, Constellation, Ajax, Zentalis and Prelude. He has received research support from and consulted for Celgene and Roche and has consulted for Syndax, Incyte, Janssen, Astellas, Morphosys and Novartis. He has received honoraria from Astra Zeneca and Novartis for invited lectures and from Gilead and Novartis for grant reviews. A.R.M. and R.H. receive funding from Dompè Pharmaceuticals. J.C., E.T., T.M., and J.Z. are employees and equity partners of IsoPlexis Corporation. M.K. is currently an employee of Imago BioSciences. No other authors report competing interests.

## STAR METHODS

### RESOURCE AVAILABILITY

#### Lead contact

Further information and requests for resources and reagents should be directed to and will be fulfilled by the Lead Contact, Ross Levine.

#### Materials availability

This study did not generate new unique reagents.

#### Data and Software Availability

De-identified patient data have been deposited to Gene Expression Omnibus (GEO). Data are publicly available as of the date of this publication. Accession numbers are listed in the key resources table. The data set contains the single-cell RNA-sequencing data shown in Figures 1 and Supplemental S1, as well as bulk RNA- and ATAC-sequencing data shown in Figure 3 and Figure S4. This paper also analyzes existing, publicly available data from Norfo et al.^38^ The accession number for this dataset is listed in the key resources table.

## EXPERIMENTAL MODEL AND SUBJECT DETAILS

### Human Subjects

Peripheral blood and splenic patient samples were provided by the Tissue Bank of the Myeloproliferative Neoplasm-Research Consortium (MPN-RC) and Memorial Sloan Kettering Cancer Center (MSKCC). Written informed consent was obtained from patients according to guidelines established by the Institutional Review Boards of Memorial Sloan Kettering Cancer Center, the Icahn School of Medicine at Mount Sinai, and each of the MPN-RC member institutions. The Institutional Review Boards of Memorial Sloan Kettering Cancer Center (MSKCC) approved sample collection and all experiments under the following protocols: 09-141, 06-107, and 16-355. All patients met the World Health Organization diagnostic criteria for myeloproliferative neoplasms (MPNs). Information about the age and gender of each patient is listed in Supp. Tables S1, S2, S3, and S4. Additional de-identified healthy bone marrow (BM) (MNCs) and purified healthy CD34^+^ cells were purchased from AllCells (Emeryville, CA). Normal splenic samples were obtained from the National Disease Research Interchange (Philadelphia, PA).

### Mouse Models

Mouse experiments were performed both at MSKCC in New York, New York (R.L.L.) and at the Department of Biomedical and Neuromotorial Sciences, Alma Mater Studiorum University in Bologna, Italy (A.R.M). All animal experiments were performed in accordance with the MSKCC Institutional Animal Care and Use Committee-approved animal protocol and by the Italian Ministry of Health (authorization 97/2021-PR; Prot. #D9997121), respectively. Animal care was performed in strict compliance with institutional guidelines, the Guide for the Care and Use of Laboratory Animals (National Academy of Sciences 1996), and the Association for Assessment and Accreditation of Laboratory Animal Care International and the Italian Law on Animal experimentation. Mice with an intact immune system were used for all experiments. All animals were drug and test naïve and not involved in previous procedures, maintained on a 12 hour light-dark cycle, and had access to water and standard chow ad libitum. Regular husbandry procedures and routine care were performed by trained veterinary staff. Animals were monitored daily for signs of disease or morbidity, failure to thrive, weight loss, open skin lesions, bleeding, infection, or fatigue and sacrificed immediately if they exhibited any signs of the above. *Cxcr2* knock-out (KO) and *Gata1^low^* mice have been described previously.^45,66^ *Cxcr2* floxed mice were crossed to *Vav-Cre* deleter lines. Age matched males and females were used as donors for *Cxcr2* KO vs. wild-type (WT) hMPL^W515L^ experiments. Female C57BL/6J mice were used as recipient mice for all *Cxcr2* KO vs. WT hMPL^W515L^ transplant experiments. Female BalbC mice were used as recipient mice for all reparixin/ruxolitinib hMPL^W515L^ *in vivo* drug trials. There was no difference observed in results obtained with females and males when analyzed separately for the primary mouse *Gata1^low^* studies.

## METHOD DETAILS

### Human cell positive selection

For patient CD34+ cell selection for single-cell RNA-Seq, single-cell cytokine, bulk RNA-Seq/ATAC-Seq, and *in vitro* culture experiments, mononuclear cells (MNC) were purified using Ficoll-Pacque (GE Healthcare Life Sciences) and CD34+ cells isolated using human CD34 MicroBeads (Miltenyi) and column filtration per the manufacturer’s protocol. For single-cell RNA-Seq and cytokine experiments, sorted cells were directly loaded into a micro-chamber array for multiplexed cytokine profiling as well as closed microwell array chip for single cell 3’ mRNA sequencing. For bulk RNA-Seq and ATAC-Seq, cells were sorted by fluorescent assisted cell sorting on the SH800S cell sorter (Sony Biotechnology) using the following antibodies: FITC anti-human lineage cocktail (Cat# 343604, BioLegend) and APC anti-human CD34 (Cat# 343509, BioLegend) and sorted directly into Trizol or PBS respectively, and ATAC library samples prepared the same day. RNA was extracted using the Direct-zol RNA microprep kit (Zymo; Cat# R2061). For megakaryocyte culture experiments, human CD41+ cells were isolated using the CD41+ selection TAC kit (Cat# 10450, StemCell Technologies).

### Single-cell RNA Sequencing

Single cell 3’ mRNA sequencing was conducted by using closed microwell arrays as previously described.^30^ In brief, isolated, CD34+ enriched cells from MPN patients and uniquely barcoded mRNA capture beads were co-isolated in individual microwell arrays followed by lysis to capture single cell mRNA. To prevent cross-contamination, fluorinated oil (Fluorinert FC-40, Cat# 86508-42-1, Sigma-Aldrich) was loaded into the device for sealing microwell arrays after introducing freeze-thaw lysis buffer. Isolated single cells were lysed by three freeze-thaw cycles and then incubated at room temperature for 1 hour to allow mRNA capture onto the capture bead. Following incubation, beads were purged out from microwell arrays after inverting the device and collected into an Eppendorf tube. Reverse transcription, library construction, and sequencing was followed previously reported protocols.^67,68^ The sequencing library was constructed by using the Nextera XT tagmentation kit (Cat# FC-131-1096, Illumina), and the HiSeq2500 instrument (Illumina) was used for a sequencing of constructed libraries using 75bp pair-end reads. Cell barcodes and unique molecular identifiers (UMIs) were generated from raw reads, and the human genome (hg19) using STAR v2.5.2b was used for alignment based on the Dropseq method.^67^ Only cells with over 10,000 reads were considered for generating the digital expression matrix.

### Single Cell mRNA Sequencing Analysis

The Seurat package (V3.1.2)^69^ in R (V3.5.0) was used to identify differentially expressed genes from 7 patients (see Supp. Table S1). For the downstream analysis, only cells with more than 100 genes and fewer than 5,000 genes as well as less than 10% mitochondrial genes were considered for analysis. Following filtration, a total of 97,412 genes in 8,144 single cells were included in the final analysis. Using the Seurat function FindVariableGenes, 2,719 variable genes were selected. After regressing cell cycle relate genes, unsupervised clustering in principal component analysis (PCA) was conducted by using the 1,000 highly variable genes from all the samples. Then, uniform manifold approximation and projection (UMAP) was used to project single cells onto a two-dimensional map with a resolution of 0.5. The SingleR package (V1.0.1)^70^ was used to annotate cell types of 8,144 single cells in R. Firstly, annotating cell lineage of each cell was followed by the authors’ instructions, which is based the data set from Blueprint Epigenomics and Endocode as a reference gens for human samples.^71,72^ The digital expression matrix was then generated for each sub-population with individual UMAP plots. Using a gene set in each cluster, functional enrichment analysis was conducted to identify the top 10 gene ontology (GO) pathways in each cluster by using the Databases for Annotation, Visualization and Integrated Discovery (DAVID).^73^ Significant correlations between genes were assessed using the R package Scran (v1.12.1).

### Single-cell multiplexed cytokine profiling

Single-cell cytokine profiling was carried out on isolated patient CD34+ cells using a previously described method.^34^ See Supp. Table S2 for a full list of patients on whom single cell cytokine analysis was performed. Briefly, a high-density antibody barcode was created using a microchannel-guided flow patterning technique as previously described. Single cell suspensions of bone marrow cells were pipetted onto the microchamber chip with individual wells pre-labelled with individual cytokine antibody barcodes. After incubation (24 hours, 37°C in 5% CO_2_), the cytokine signals were then developed using an immuno-sandwich ELISA assay and scanned with a GenePix 4200A (Axon) microarray scanner to collect fluorescent signals. Signals were then processed with GenePix software and Excel macro to acquire average fluorescent signals from each microchamber for all bars in each antibody barcode. Single-cell data were gated on the basis of the background signals to distinguish between cytokine producers and nonproducers. The data were further transformed and normalized to perform multidimensional data analysis as previously described.^34^

### Data Processing and Analysis of Single-Cell Cytokine Assays

For all single-cell secretomic analysis, custom-built algorithms in Excel, MATLAB, and R-packages were used to process, analyze, and visualize the single-cell cytokine profiles. Briefly, the predetermined cell numbers of the microchambers were normalized to fluorescence intensity values for each measured protein from corresponding microchambers. Background signals from empty microchambers were used to draw gates for distinguishing cytokine-producing and non-producing cells. The gating threshold was calculated by (average intensity of all empty microchambers for a given cytokine) + 3 × (standard deviation of all empty microchambers for a given cytokine). Any cells with fluorescence intensity of a specific cytokine below the gating threshold were considered non-producers, and their cytokine intensity values were converted to zero. Any cells with fluorescence intensity above the gating threshold were considered cytokine-producers and given the cytokine intensity value of: (the measured fluorescence intensity − the average fluorescence intensity of all zero-cell microchambers for a given cytokine). The gated and background-subtracted cytokine intensity values were then log-transformed. Protein levels were normalized by the average intensity and the standard deviation to eliminate variability in detection sensitivity and secretion profile of assayed proteins. The number of all cells above the gates and the sum of fluorescence intensity of a given cytokine were measured to calculate the frequency of cytokine producers and the average cytokine secretion per single cell.

### *In vitro* culture experiments

For liquid culture experiments, CD34+ cells from MF patients were isolated as described above and cultured in StemSpan serum free medium (SFEM; cat# 09605, StemCell Technologies) with stem cell factor (SCF) (Cat# 255-SC, R&DSystem), thrombopoietin (TPO; Cat# 288-TP, R&DSystem), FMS-like tyrosine kinase 3 ligand (FLT3L; Cat# 308-FKN, R&DSystem) and interleukin-3 (IL-3; Cat# 203-IL, R&DSystem) at 20 ng/ml of each, to which various doses of CXCL8 (0, 50, and 100 ng/ml, respectively) (Cat# 208-IL, R&DSystem) were added. Cell viability was determined using cellometer Auto 2000 (Nexcelom Bioscience). The proportion of CD34, CD41 and CD33 cells was measured by flow cytometry. Megakaryocyte (MK) culture experiments were carried out using previously described methods.^46^ Briefly, isolated CD34^+^ cells were cultured in StemSpan SFEM with 100 ng/ml of SCF and 50ng/ml of TPO for 7 days and then harvested and washed with phosphate buffered saline (PBS; Cat# 21-040-CM, Corning) and cultured once more in SFEM with 50 ng/ml of TPO alone for an additional 7 days to generate a cell population enriched for both MK progenitor (CD34^+^/CD41^+^) and mature (CD34^-^/CD41^+^ and CD41+/CD42+ cells) MKs. At time of analysis, cells were counted using cellometer Auto 2000 (Nexcelom Bioscience) and samples were washed with MACS buffer (Cat # 130-092-987, Miltenyi Biotec) and stained for the following antibodies for fluorescence-assisted cell sorting: CD34 (Cat# 555824, BDBiosciences), CD33 (Cat# 555450, BDBiosciences), CD41 (Cat# 555824, BDBiosciences), CD45 (Cat# 555485, BDBiosciences), CXCR1 (Cat# 555939, BDBiosciences) or CXCR2 (Cat# 320706, BioLegend) and acquired and analyzed on a FACSCaliber analyzer (BDBiosciences).

### Methylcellulose Assays

CD34+ cells isolated from MF patients were plated at a density of 500 cells/replicate in 30-mm dishes containing 1 mL of StemSpan SFEM with 1.1% methylcellulose (Cat# H4230, StemCell Technologies), to which SCF, TPO, FLT3L, granulocyte macrophage–colony stimulating factor (Cat # 215-GM, R&DSystem), IL-3, and erythropoietin (EPO; Cat # 287-TC, R&DSystem) were added, with or without CXCL8. For reparixin/ladarixin studies, DMSO was added to control wells. All experiments were performed in duplicate. Colony-forming unit-granulocyte-macrophage (CFU-GM) and CFU-GEMM colonies were enumerated after 14 days incubation and normalized as a ratio to the DMSO control for each patient. For JAK2^V617F^ variant allele fraction analysis, individual colonies were randomly plucked, and JAK2^V617F^ was assayed using a previously-published nested allele-specific polymerase chain reaction (PCR).^74^ Briefly, a 521-bp DNA fragment containing the V617F mutation site was amplified from 2uL of gDNA using primers P1 5′-GATCTCCATATTCCAGGCTTACACA-3′ and P1r 5′-TATTGTTTGGGCATTGTAACCTTCT-3′. After 35 cycles consisting of 30s at 95°C, 30s at 60°C, and 30s at 72°C, 2μL PCR products were further amplified by nested and allele-specific primers P2 5′-CCTCAGAACGTTGATGGCA-3′, P2r 5′-ATTGCTTTCCTTTTTCACAAGA-3′, Pnf 5′-AGCATTTGGTTTTAAATTATGGAGTATATG-3′, and Pmr 5′-GTTTTACTTACTCTCGTCTCCACAAAA-3′, following 35 cycles of 95°C for 30s, 59°C for 25s, and 72°C for 25s. The final PCR products were analyzed on 2.5% agarose gels. The nested PCR product had a size of 453bp. A 279-bp product indicated allele-specific JAK2^V617F^ positive whereas a 229-bp product denoted allele-specific wild product. Colonies classified as homozygous for JAK2^V617F^ contained only the 279-bp band, whereas heterozygous colonies were identified based on the presence of both the 279-bp and 229-bp bands.

### Isolation of RNA and qRT-PCR for *in vitro* culture studies

Total RNA was extracted from CD34+ and CD41+ cell populations using the RNeasy kit (QIAGEN, Valencia, CA). Complementary DNA was reverse transcribed using EcoDry Premix kit (TaKaRa, Mountain View, CA). The *CXCL8* (GeneGlobe ID: PPH00568A-200), *VEGF* (GeneGlobe ID:PPH00251C-200), *TGF-β1* (GeneGlobe ID: PPH00508-200), *IL-6* (GeneGlobe ID: PPH00560C-200), and *GAPDH* (GeneGlobe ID: PPH00150F-200) genes were evaluated by quantitative reverse-transcription (qRT) PCR, using SYBR Green RT2 qPCR Master mixes from QIAGEN and the RealPlex thermocycler (ThermoFisher Scientific, Fairlawn, NJ). All primers for qRT-PCR were purchased from GeneGlobe.

### CXCL8 Immunohistochemistry

For primary MF and healthy bony marrow and spleen samples, sections of formalin-fixed and paraffin-embedded marrow biopsy and spleen samples were baked and de-paraffinized. Immuno-histochemical staining for CXCL8 (Cat# ab106350, Abcam) was performed using the Bond III auto-stainer (Leica Microsystems). The degree of fluorescence intensity was assessed using MetaMorph Microscopy Automation and Image Analysis Software (Molecular Devices).

### Patient Plasma Cytokine Analysis

CXCL8 plasma levels in normal and MF patients were evaluated using the ELISA Human CXCL8 Quantikine ELISA Kit (Cat# D8000C, R&DSystem). Luminex assays for murine serum were carried out using the FlexMAP 3D multiplexing platform (Luminex xMAP system). The Millipore Mouse Cytokine 32-plex kit was used to measure the serum concentration of 32 cytokines (Cat# MCYTMAG-70K-PX32, Millipore). xPONENT (Luminex) and Milliplex Analyst Software (Millipore) was applied to convert mean fluorescent intensities (MFI) values into molecular concentrations by the use of a standard curve (5-parameter logistic fitting method).

### RNA-seq Analysis

Fastq files containing RNA-seq paired-end reads were trimmed for both base quality (q <=15) and Illumina adaptor sequences using TrimGalore v0.4.5 (https://github.com/FelixKrueger/TrimGalore; Krueger F, Trimgalore, 2021). STAR v2.6.0a^75^ was used to align the RNA-seq fastq files to the reference genome (hg19) and to count the number of reads mapping to each gene. Duplicates were removed using Picard MarkDuplicates v2.16 (https://broadinstitute.github.io/picard/command-line-overview.html#MarkDuplicates). Gencode annotations of the hg19 human genome was used for the genomic location of transcription units.^76^ Genome-wide transcript counting was then performed by featureCounts v1.6.3.^77^ We used the median of ratios method of normalization from DESeq2^78^ to transform the data into a normalized read count matrix. Differential expression between the different groups (CXCL8 high vs CXCL8 low secretors in the myelofibrosis patient cohort) was performed through the use of DESeq2 1.22.2^78^ with a fold change cutoff of ± 2 and a FDR of 1%. Motif signatures were obtained using the ‘de novo’ approach in Homer v4.11^79^ and then matched to the default ‘known TF’ database. Gene set enrichment analysis was performed using GSEA v3.0^80^ with mSigDB (v6.0) pathway database to estimate the overrepresented gene set pathways in the differential expression analysis.

### Protein-protein interaction (PPI) network analysis

We visualized the protein interaction network of the differentially upregulated genes with fold change of >= 4 and FDR of <= 1% from RNA-seq that interacted with each other using the STRING database v11.5.^81^ For visualization, the STRING protein query network at a confidence of >= 0.40 was imported to Cytoscape v3.8.2.^82^ The degree and betweenness centrality was calculated for each node using NetworkAnalyzer v4.4.6.^83^ The STRING network was then filtered for nodes with degree >= 5 and betweenness centrality score >=0.005 to visualize a concise sub-network module. STRING functional enrichment was performed with Gene Ontology (GO), InterPro, Kyoto Encyclopedia of Genes and Genomes (KEGG) Pathways, and PFAM databases.^81^

### ATAC-seq Analysis

Fastq files containing ATAC-seq paired-end reads were trimmed for both base quality (q <=15) and Illumina adaptor sequences using TrimGalore v0.4.5. Trimmed reads were mapped to the hg19 human genome using Bowtie2 v2.3.4.1.^84^ Duplicates were removed using Picard MarkDuplicates v2.16. Peak calling was performed using MACS2 v2.1.2 against standard input as the control (fold change > 2 and p value < 0.001).^85^ The peaks from all samples were then merged within a 500 bp window to create a full peak atlas. Raw read counts were tabulated over this peak atlas using featureCounts v1.6.3.^77^ Peaks were annotated using genomic distance, with a gene assigned to a peak if it was within 50 kb up- or down- stream of the gene start or end site. The raw read counts in the peak atlas were normalized using the median of ratios normalization method in DESeq2 v1.22.2 to obtain a normalized read count matrix.^78^ Differential accessibility of these peaks was then calculated in the comparison of CXCL8 high vs low secretors in the myelofibrosis patient cohort using DESeq2 with a fold change cutoff of ± 2 and a FDR of 1%.

### RNA- and ATAC-Sequencing Integration Analysis

For the integrated analysis of RNA and ATAC-seq, the RNA-seq samples were subsetted to those that had the respective ATAC-seq data available (See Supp. Table S3). The differential expression of this subset of samples with available ATAC-seq data was calculated in the same comparison of CXCL8 high vs low secretors as in the ATAC-seq differential analysis in the myelofibrosis patient cohort using DESeq2 v1.22.2. Peaks were annotated with an associated gene using genomic distance, with a gene assigned to a peak if it was within 50 kb up- or down- stream of the gene start or end site. We visualized the RNA differential expression and ATAC differential accessibility associated with a gene in a waterfall plot. The top 50 up-regulated and down-regulated genes were plotted with FDR <= 2.5% for RNA-seq differentially expressed genes and nominal p-value <= 2.5% for those associated ATAC-seq differential accessibility regions in closest proximity to the gene. The log2 fold change of the gene expression of the top differentially expressed and differentially accessible genes was plotted as a waterfall bar plot and the associated peaks were plotted as colored circles whose size and color and distance along the x-axis represent the log2 fold change in differential accessibility of the gene.

### Microarray Analysis (public dataset)

GSE53482 from Gene Expression Omnibus (GEO) consists of the gene expression profile of CD34+ cells from 42 PMF patients and 31 healthy donors performed on the Affymetrix Human Genome U219 Array.^38^ The public myelofibrosis microarray data CEL files were downloaded and processed using the oligo v1.54.1 package in R v4.0.3.^86^ Robust Multi-array Average (RMA) was used to normalize the raw microarray expression values.^87^ The probes were annotated using their gene symbol names with annotate v1.68, hgu219.db v3.2.3, and hgu219cdf v2.18 R packages. The patient cohort was then divided into two groups of patients based on their CXCL8 expression: specifically CXCL8 high which were the patients with CXCL8 expression greater than the 80% quantile and CXCL8 low which were the patients with CXCL8 expression less than the 20% quantile. The differential expression of genes was then calculated in the comparison of CXCL8 high (>= 80 % quantile) vs low (<= 20% quantile) expression in the myelofibrosis patient cohort using the limma v3.46 package with a fold change cutoff of ± 1.5 and a FDR of 5%.^88^ Gene set enrichment analysis was performed using GSEA v3.0 with mSigDB (v6.0) pathway database to estimate the enriched gene set pathways. We plotted the heatmap of all the significantly differentially expressed genes using the complement of the pearson correlation as the measure of distance with Ward’s method for hierarchical clustering, a method that minimizes the variance or the sum of squared errors between samples to find the centroids of the data to visualize the differentially expressed genes in distinct gene clusters.^89^

### hMPL^W515L^ Transplantation and *in vivo* inhibitor experiments

MPL^W515L^ bone marrow transplantation experiments were performed as described previously.^39^ For genetic deletion studies, pre-stimulated, c-KIT enriched bone marrow cells from *Cxcr2^f/f^;VavCre^+^* or *Cxcr2^f/f^;VavCre^-^* wild-type (WT) donor mice were subjected to one round of co-sedimentation with viral supernatant containing MSCV-*hMpl*^W515L^-IRES-GFP plasmid over retronectin-coated plates (Takeda). A total of 4-5 × 10^5^ cells (∼25%–40% GFP-positive, MPL^W515L^-expressing cells) were then injected into the tail veins of lethally irradiated congenic CD45.1 mice (purchased from Jackson Laboratories) along with 50,000 WT CD45.1 support marrow. For *in vivo* drug trial experiments, BalbC donor and recipient mice were used for hMPL^W515L^ transfection experiments and were prepared in a similar fashion to that as above. Approximately 10-14 days post-transplant, at first signs of disease, mice were randomized to begin 21 days of treatment with the JAK1/2 inhibitor ruxolitinib (60 mg/kg P.O. twice daily), reparixin (60mg/kg SubQ twice daily), combination therapy, or vehicle. Mice were ranked on the basis of peripheral WBC count and assigned to treatment groups to achieve congruent WBC profiles. Investigators were not blinded to the identity of mice or samples. Blood sampling was performed by submandibular bleeding using a 5mm lancet (MEDipoint Inc) and automated peripheral blood counts obtained using a ProCyte Dx (IDEXX Laboratories) according to standard manufacturer’s instruction. Timed sacrifice studies were completed at 7-8 weeks post-transplant. At time of sacrifice, bone marrow was harvested and spleens and livers were removed and weighed, and single-cell suspensions were prepared for subsequent cell staining and fractionation, flow cytometry, Western blot analysis, or histopathologic analysis. Peripheral blood serum was collected for cytokine analysis by incubating whole blood in Eppendorf tubes for 30 minutes followed by centrifugation at 2000rpm for 10 minutes and collection of supernatant.

### Histopathologic analysis

Tibia and spleen samples were fixed in 4% paraformaldehyde >24 hours and then embedded in paraffin. Paraffin sections were cut on a rotary microtome (Mikrom International AG), mounted on microscope slides (Thermo Scientific), and air-dried at 37°C overnight. After drying, tissue section slides were processed either automatically for hematoxylin and eosin (H&E) staining (COT20 stainer, Medite) or manually for reticulin staining. Pictures from H&E and reticulin stained slides were using BX53 Olympus microscope and DP74 mounted camera (original magnification 200x) and analyzed with Olympus Cellsens software. Murine tissue sections were formally evaluated by a hematopathologist (W. Xiao), including reticulin scoring. For analysis of tissue from *in vivo* inhibitor studies, samples were blinded.

### Antibodies, FACS, and western blot analysis

Following single cell preparation, blood, whole marrow, or spleen mononuclear cells were stained with a lineage cocktail comprised of antibodies targeting CD3, B220, CD41, Gr-1, CD11b, and Ter119. Cells were also stained with antibodies against c-Kit, Sca-1, FcγRII/III, and CD34. Cell populations were analyzed using a Fortessa Flow Cytometer (BD). All FACS antibodies were purchased from BD, BioLegend, or eBioscience. We used the following antibodies: c-Kit (2B8), Sca-1 (D7), Mac-1/CD11b (M1/70), Gr-1 (RB6-8C5), Ter-119 (Ter-119), CD34 (RAM34), FcγRII/III (2.4G2), CD45.1 (A20), CD45.2 (104), CD45R/B220 (RA3-6B2), CD71 (R17217), CD41 (MWReg30), CD3 (17A2), CXCR2 (SA044G4). For Western blot analysis, whole-cell protein extracts were prepared with RIPA buffer containing a protease inhibitor cocktail (Thermo Scientific). Proteins were separated by NuPAGE 4-12% Bis-Tris Gel (Thermo Scientific) and transferred to a nitrocellulose membrane using iBlot 2NC regular stacks transfer system (Invitrogen). The following antibodies were used for western blot analysis: S100a8 (Cat# 47310S, Cell Signaling), S100a9 (Cat# 73425S, Cell Signaling), pSTAT3 (Cat# 9145S, Cell Signaling), STAT3 (Cat# 4904S, Cell Signaling), pERK (Cat# 4370S, Cell Signaling), ERK (Cat# 4695S, Cell Signaling), and H3 histone (Cat# 9715S, Cell Signaling).

### *Gata1^low^* inhibitor studies

Age-matched eight-month-old *Gata1^low^* mice (N=8 males and N=8 females) were anesthetized with 2-to-3% isoflurane and implanted subcutaneously by making a mid-scapular incision with ALZET® Osmotic Pumps (model 2002) pre-filled with 200 μL of vehicle (sterile saline) or reparixin (7.5mg/h/Kg in sterile saline) as described by manufacturer’s instruction. The concentration of reparixin for subcutaneous delivery was calculated by dividing the total dose administered (per hour) by the pumping rate of the pump (see: https://www.alzet.com/formulating-thesolution/). Following 20 days of treatment, three mice per experimental group were weighed, bled for blood parameters, and sacrificed for organ histopathology observations. BM and spleen tissues were fixed in 10% phosphate-buffered formalin. Paraffin-embedded tissue was cut into consecutive 2.5–3μm sections and stained either with Gomori silver (Cat# 04-040801, Bio-Optica) or Reticulin (Cat# 04-040802, Bio-Optica), which both detect reticulin fibers. BM sections were immunostained with Cxcl1 (Cat# ab86436, Abcam), Cxcr1 (Cat# GTX100389, Genetex) and Cxcr2 (Cat# ab14935, Abcam) antibodies. Images were acquired with an optical microscope (Eclipse E600, Nikon) equipped with the “33” Series USB 3.0 Camera (Cat# DFK 33UX264, Imaging Source). Images were processed and the intensity of the immunostaining quantified with the software ImageJ (version 1.52t; National Institutes of Health) and expressed as a percent of area stained above threshold.

### Splenic stromal/fibroblast cells and endothelial cells preparation

Stromal cells were isolated from normal spleen donors obtained from the National Disease Research Interchange (healthy donors who required splenectomy after a traumatic injury). Tissue was cut into small pieces and digested using collagenase (Cat# 501004980, Thermo Scientific) and single cells washed with PBS. To generate fibroblasts, digested splenic cells were then cultured in MEM-α with 5% human platelet lysates (PLTMax; Cat# SCM142, Millipore Sigma) at a density of 2-5 × 10^4^ cells/mL in culture flasks for 24 hours. The cells that remained in suspension were removed by completely changing the medium. The medium was changed twice weekly by replacing half of the medium with fresh MEM-α. To generate endothelial cells, the digested splenic cells were cultured in endothelial cell growth medium EGM-2 (Cat# CC-3162, Lonza) at a density of 2-5 × 10^4^/mL in fibronectin-coated culture flasks for 24 hours, then the suspension cells were removed by completely changing the medium. The medium was changed twice weekly by replacing half of the medium with fresh EBM-2.

For co-culture experiments, 5 × 10^4^ healthy donor or MF megakaryocyte (MK) enriched cells were directly seeded onto 12-well plates over top mesenchymal stromal cells (MSCs) at 50% confluency. To evaluate the effects of CXCL8 enumeration in response to CXCR1/2 inhibitor therapy, 10uM of Ladarixin (Dompè) was added (or DMSO as control). After 3 days of co-cultivation, the media were collected and analyzed for secretion of CXCL8 and VEGF using respective ELISA kits (CXCL8: Cat# D8000C, R&DSystem; VEGF: Cat# DVE00, R&DSystem); the results were then normalized by cell number. Both suspension and adherent cells were collected and stained with anti-hCD45 (Cat# 555485, BDBiosciences) to identify the hematopoietic cells from adherent cells.

## QUANTIFICATION AND STATISTICAL ANALYSIS

Colony formation assays and liquid culture experiments using human cells shown in this study were performed in duplicate. For retroviral MPL^W515L^ BMT experiments, blood count analysis and organ weights of vehicle- and drug-treated mice were recorded as indicated in figure legends. Donor and recipient mice for the retroviral MPL^W515L^ BMT experiments were used at a ratio of 1:5. The CXCR2 genetic knock-out hMPL^W515L^ studies were performed for at least 3 independent experiments. The *in vivo* experiments were designed to use the minimum number of animals required. Pathologic sections were performed for a total of N=6 (vehicle) and N=6 (drug-treated) mice to assess the effect of reparixin +/- JAK inhibition *in vivo*. Survival experiments were performed twice with N=6-12 per arm. The number of animals, cells, and experimental replication can be found in the respective figure legends. No statistical methods were utilized to determine sample size. The Student’s t-test (unpaired, two-tailed) was used to compare the mean of two groups. Normality tests were used to test the assumption of normal distribution. Data were analyzed and plotted using GraphPad Prism 7 software. Graphs represent mean values ± standard error of the mean (SEM) for most *in vivo* data and mean ± standard deviation (SD) for most *in vitro* data. Kaplan-Meier survival analysis and long-rank test was used to compare survival outcomes. For multiple comparisons, one-way or two-way analysis of variance (ANOVA) with post hoc Bonferroni correction was used. Statistical significance was established at P< 0.05.

---

**Supplemental Table 1.** Selected patients for single cell transcriptional (scRNA-Seq) analysis.

| ID | Diagnosis | Age | Sex | Driver mutation | Grade reticulin fibrosis | Sequenced cell number (after filtration) | Sample Origin |
| --- | --- | --- | --- | --- | --- | --- | --- |
| MF1 | Primary myelofibrosis | 76 | F | JAK2<sup>V617F</sup> | 3+/3 | 801 | Patient Blood |
| PV1 | Polycythemia vera | 59 | F | JAK2<sup>V617F</sup> | 0-1+/3 | 892 | Patient Blood |
| PV2 | Polycythemia vera | 41 | M | JAK2<sup>V617F</sup> | 1-2+/3 | 772 | Patient Blood |
| PV3 | Polycythemia vera | 46 | M | JAK2<sup>V617F</sup> | 0/3 | 700 | Patient Blood |
| ET1 | Essential thrombocythemia | 74 | F | JAK2<sup>V617F</sup> | 1+/3 | 532 | Patient Blood |
| ET2 | Essential thrombocythemia | 40 | M | CALR | 0-1+/3 | 505 | Patient Blood |
| ET3 | Essential thrombocythemia | 73 | M | JAK2<sup>V617F</sup> | 2+/3 | 997 | Patient Blood |

MF: myelofibrosis; PV: polycythemia vera; ET: essential thrombocythemia

**Supplemental Figure 1:**
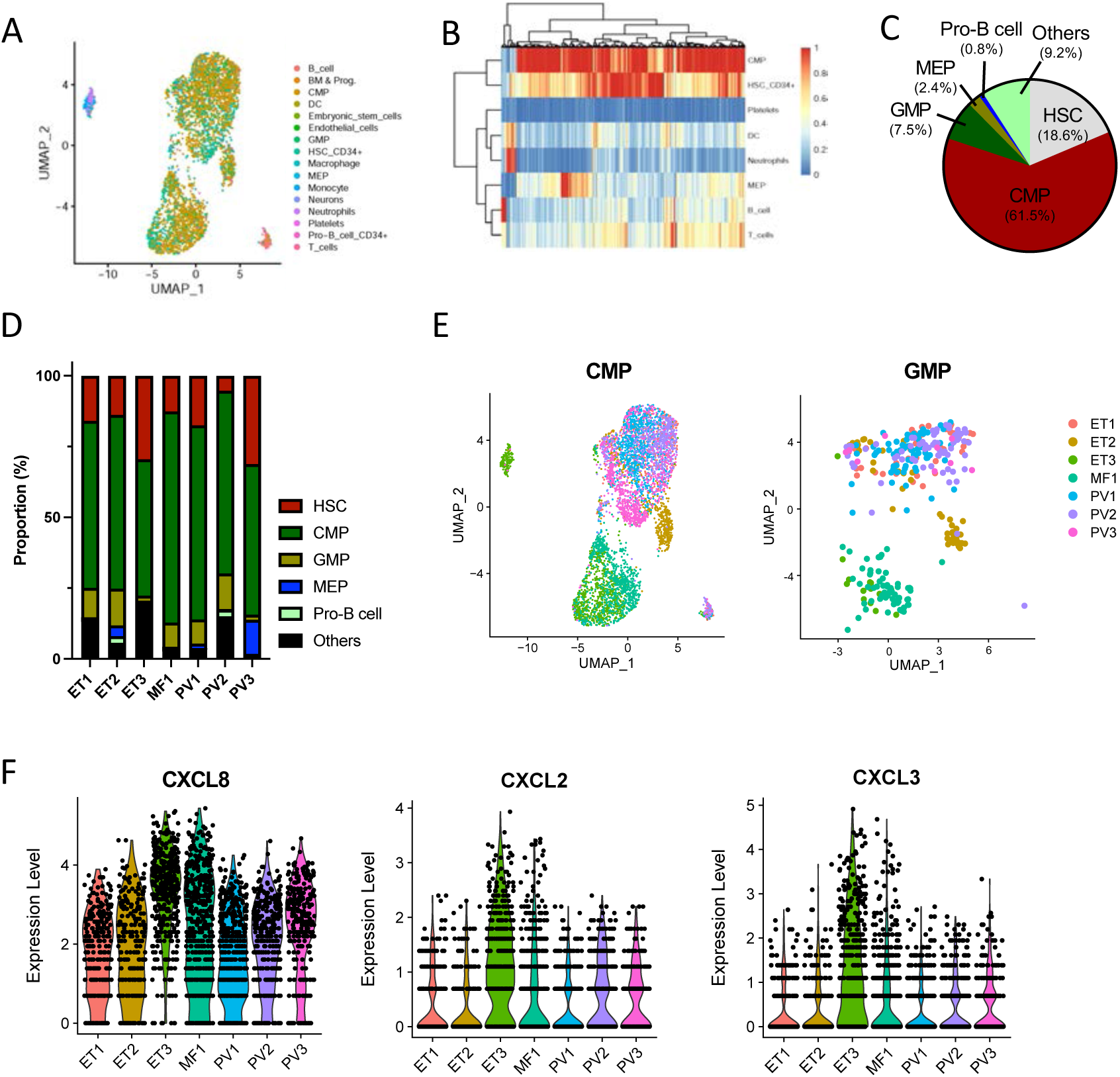
Single-cell RNA-Seq (scRNA-Seq) identifies enrichment in CXCR2 signaling mediators. **(A)** Uniform manifold approximation and projection (UMAP) demonstrating clustering of detectable cell populations based on single-cell transcriptional profiling. Each dot represents a single cell with color corresponding to cell type (CMP, common myeloid progenitor; DC, dendritic cell; GMP, granulocyte monocyte progenitor; HSC, hematopoietic stem cell; MEP, megakaryocyte erythroid progenitor). **(B)** Hierarchical clustering of detectable cell populations based on transcriptional profiling. Each hashmark represents a single cell. Colors represent normalized scores. **(C)** Pie chart depicting percent frequency of classifiable hematopoietic cell populations identified by scRNA-Seq across all 7 patients.(CMP, common myeloid progenitor; GMP, granulocyte monocyte progenitor; HSC, hematopoietic stem cell; MEP, megakaryocyte erythroid progenitor). **(D)** Percent frequency of detectable cell populations identified by scRNA-Seq per patient. **(E)** UMAPs of common myeloid progenitor (CMP; left panel) and granulocyte-monocyte progenitor (GMP; right panel) populations colored by patient (ET, essential thrombocythemia; PV, polycythemia vera; MF, myelofibrosis). Each dot represents a single cell. **(F)** Violin plots showing relative expression of CXCR2-mediated cytokines *CXCL8* (left panel), *CXCL2* (middle panel), and *CXCL3* (right panel) in CMPs of individual MPN patients by scRNA-Seq. Each dot represents a single cell.

**Supplemental Table 2.**
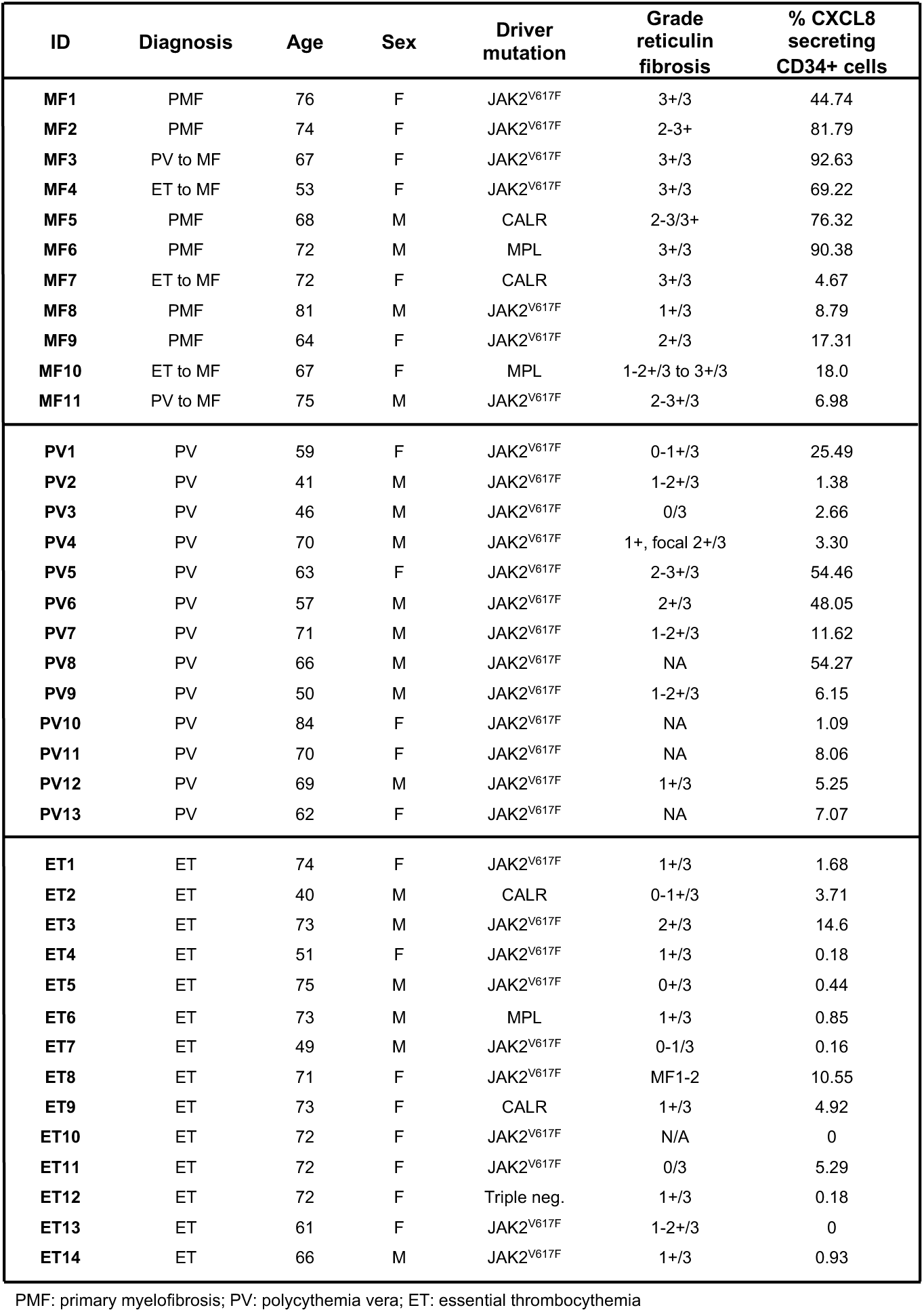
Selected patients for single-cell cytokine analysis.

| ID | Diagnosis | Age | Sex | Driver mutation | Grade reticulin fibrosis | % CXCL8 secreting CD34+ cells |
| --- | --- | --- | --- | --- | --- | --- |
| MF1 | PMF | 76 | F | JAK2<sup>V617F</sup> | 3+/3 | 44.74 |
| MF2 | PMF | 74 | F | JAK2<sup>V617F</sup> | 2-3+ | 81.79 |
| MF3 | PV to MF | 67 | F | JAK2<sup>V617F</sup> | 3+/3 | 92.63 |
| MF4 | ET to MF | 53 | F | JAK2<sup>V617F</sup> | 3+/3 | 69.22 |
| MF5 | PMF | 68 | M | CALR | 2-3/3+ | 76.32 |
| MF6 | PMF | 72 | M | MPL | 3+/3 | 90.38 |
| MF7 | ET to MF | 72 | F | CALR | 3+/3 | 4.67 |
| MF8 | PMF | 81 | M | JAK2<sup>V617F</sup> | 1+/3 | 8.79 |
| MF9 | PMF | 64 | F | JAK2<sup>V617F</sup> | 2+/3 | 17.31 |
| MF10 | ET to MF | 67 | F | MPL | 1-2+/3 to 3+/3 | 18.0 |
| MF11 | PV to MF | 75 | M | JAK2<sup>V617F</sup> | 2-3+/3 | 6.98 |
| PV1 | PV | 59 | F | JAK2<sup>V617F</sup> | 0-1+/3 | 25.49 |
| PV2 | PV | 41 | M | JAK2<sup>V617F</sup> | 1-2+/3 | 1.38 |
| PV3 | PV | 46 | M | JAK2<sup>V617F</sup> | 0/3 | 2.66 |
| PV4 | PV | 70 | M | JAK2<sup>V617F</sup> | 1+, focal 2+/3 | 3.30 |
| PV5 | PV | 63 | F | JAK2<sup>V617F</sup> | 2-3+/3 | 54.46 |
| PV6 | PV | 57 | M | JAK2<sup>V617F</sup> | 2+/3 | 48.05 |
| PV7 | PV | 71 | M | JAK2<sup>V617F</sup> | 1-2+/3 | 11.62 |
| PV8 | PV | 66 | M | JAK2<sup>V617F</sup> | NA | 54.27 |
| PV9 | PV | 50 | M | JAK2<sup>V617F</sup> | 1-2+/3 | 6.15 |
| PV10 | PV | 84 | F | JAK2<sup>V617F</sup> | NA | 1.09 |
| PV11 | PV | 70 | F | JAK2<sup>V617F</sup> | NA | 8.06 |
| PV12 | PV | 69 | M | JAK2<sup>V617F</sup> | 1+/3 | 5.25 |
| PV13 | PV | 62 | F | JAK2<sup>V617F</sup> | NA | 7.07 |
| ET1 | ET | 74 | F | JAK2<sup>V617F</sup> | 1+/3 | 1.68 |
| ET2 | ET | 40 | M | CALR | 0-1+/3 | 3.71 |
| ET3 | ET | 73 | M | JAK2<sup>V617F</sup> | 2+/3 | 14.6 |
| ET4 | ET | 51 | F | JAK2<sup>V617F</sup> | 1+/3 | 0.18 |
| ET5 | ET | 75 | M | JAK2<sup>V617F</sup> | 0+/3 | 0.44 |
| ET6 | ET | 73 | M | MPL | 1+/3 | 0.85 |
| ET7 | ET | 49 | M | JAK2<sup>V617F</sup> | 0-1/3 | 0.16 |
| ET8 | ET | 71 | F | JAK2<sup>V617F</sup> | MF1-2 | 10.55 |
| ET9 | ET | 73 | F | CALR | 1+/3 | 4.92 |
| ET10 | ET | 72 | F | JAK2<sup>V617F</sup> | N/A | 0 |
| ET11 | ET | 72 | F | JAK2<sup>V617F</sup> | 0/3 | 5.29 |
| ET12 | ET | 72 | F | Triple neg. | 1+/3 | 0.18 |
| ET13 | ET | 61 | F | JAK2<sup>V617F</sup> | 1-2+/3 | 0 |
| ET14 | ET | 66 | M | JAK2<sup>V617F</sup> | 1+/3 | 0.93 |

PMF: primary myelofibrosis; PV: polycythemia vera; ET: essential thrombocythemia

**Supplemental Figure 2:**
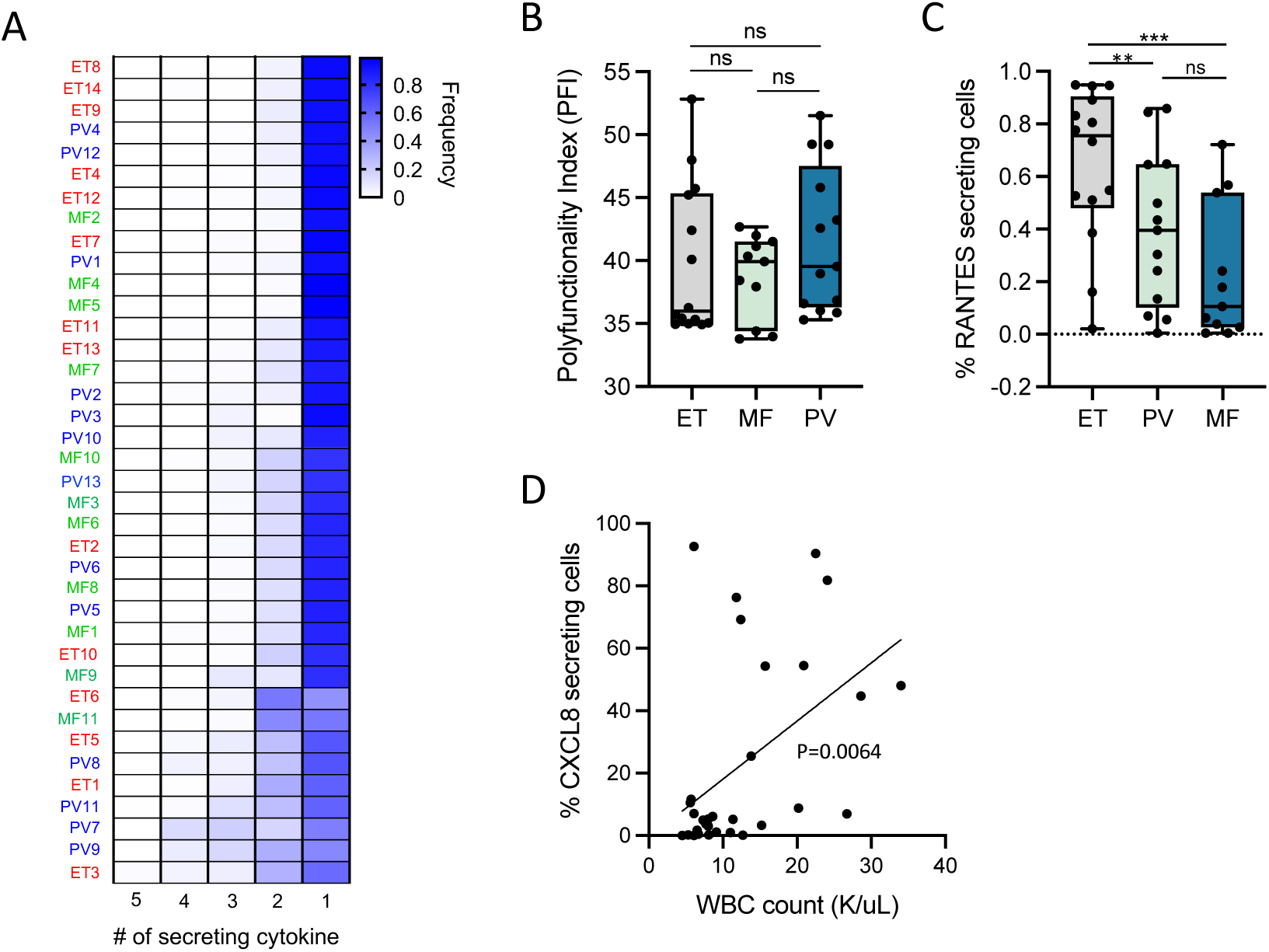
CXCL8 secretion by MF CD34+ cells correlates with clinical parameters, and MF CD34+ cells display sensitivity to CXCL8 *in vitro*. **(A)** Heatmap showing fraction of cells secreting single or >1 combined cytokines as detected by single-cell cytokine analysis for each individual patient evaluated (ET, essential thrombocythemia; PV, polycythemia vera; MF, myelofibrosis). Colors represent frequency among total cells (0% white; 100% dark blue). **(B)** Polyfunctionality index (PFI), or the fraction of multiple-secreting cytokine cell populations detected by single-cell cytokine arrays based on clinical MPN sub-type (EV, PV, or MF). ns=non-significant. **(C)** Box plots depicting correlation between MPN sub-type and percent fraction of RANTES-only secreting cells as detected by single-cell cytokine analysis. **p<0.01. ***p<0.001. ns=non-significant. **(D)** Linear correlation analysis of percent fraction of CXCL8-only secreting CD34+ cells detected by single-cell cytokine analysis and total white blood cell (WBC) count at time of sample collection.

**Supplemental Figure 3:**
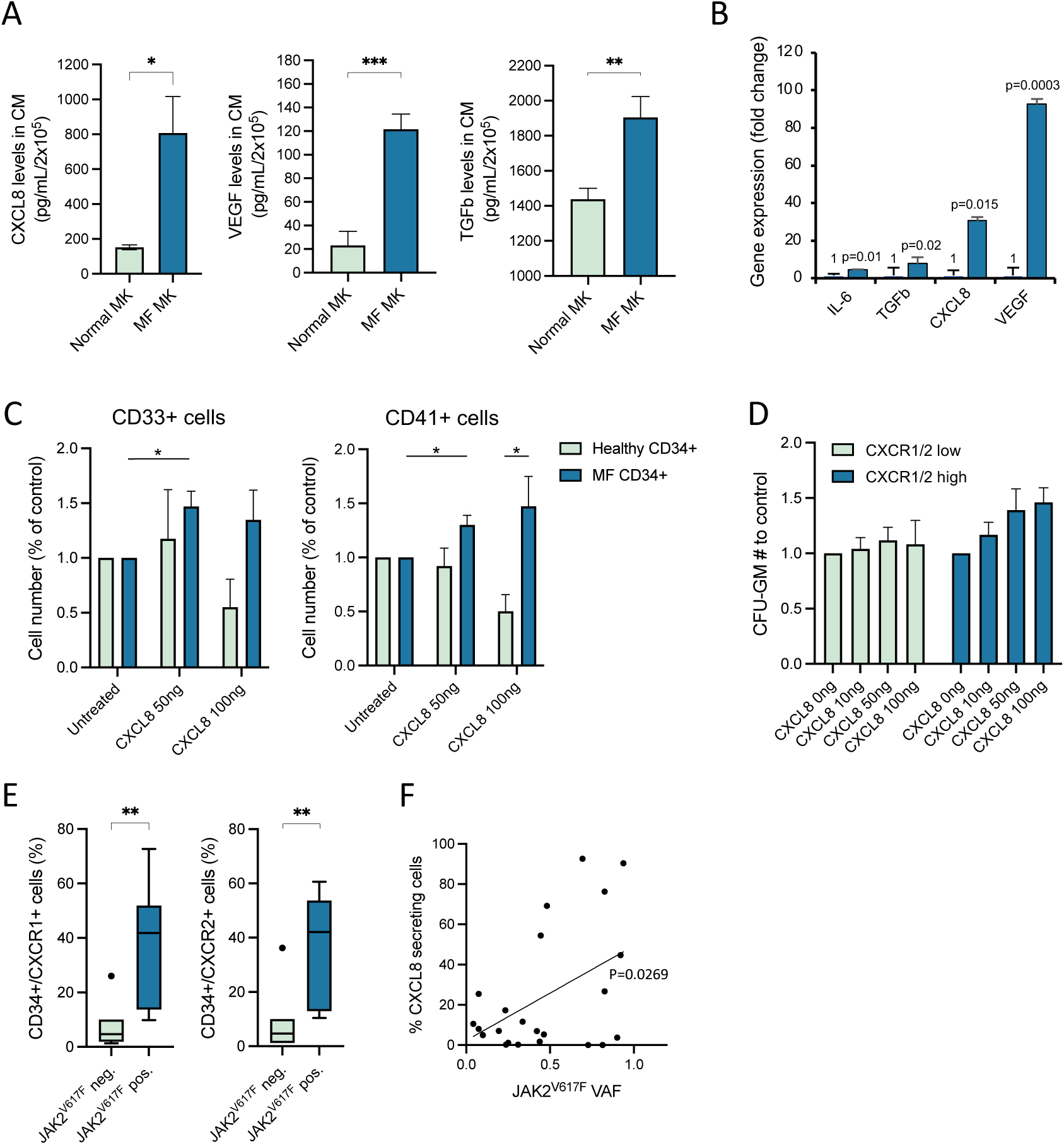
Cytokine secretion, including CXCL8, is enhanced in MF megakaryocytes (MKs) in comparison to healthy controls. **(A)** Total levels of CXCL8 (left panel), VEGF (middle), and TGFβ (right) detected in conditioned media (CM) elicited from cultured healthy donor (light blue) vs. MF (dark blue) megakaryocytes (MK). *p<0.05. **p<0.01. ***p<0.001. Data shown represent mean ± SD. **(B)** Fold change of pro-inflammatory cytokine IL-6, TGFβ, CXCL8, and TGFβ gene expression by quantitative PCR from isolated MF MKs (dark blue) compared to healthy controls (light blue). P-values as indicated. **(C)** Ratio of total CD33+ cell output (left panel) and CD41+ cell output (right panel) relative to untreated of cultured healthy donor (light blue) vs. MF (dark blue) CD34+ cells in response to exogenous CXCL8 as detected by flow cytometry. *p<0.05. Data shown represent mean ± SD. **(D)** Ratio of granulocyte-macrophage progenitor (CFU-GM) colony output to control of cultured MF CD34+ cells in methylcellulose in response to increasing doses of exogenous CXCL8 stratified by degree of CXCR1/2 surface expression (“low:” <10% of cells; “high:” ≥10% of cells) **(E)** Percentage of total MF CD34+ cells expressing CXCR1 (left panel) or CXCR2 (right panel) as detected by flow cytometry based on JAK2^V617F^ mutational status. **p-value <0.01. **(F)** Linear correlation analysis of percent fraction of CXCL8-only secreting CD34+ cells detected by single-cell cytokine analysis and JAK2^V617F^ variant allele fraction (VAF).

**Supplemental Table 3.** Selected patients for bulk RNA-Seq (+ ATAC-Seq) analysis.

| ID | Diagnosis | Age | Sex | Driver mutation | JAK2<sup>V617F</sup> VAF | Grade reticulin fibrosis | % CXCL8 secreting CD34+ cells | Sample Origin |
| --- | --- | --- | --- | --- | --- | --- | --- | --- |
| **CXCL8 secretor** |  |  |  |  |  |  |  |  |
| **MF2\*** | PMF | 74 | F | JAK2<sup>V617F</sup> | N/A | 2-3+ | 81.79% | Patient Blood |
| **MF3\*** | PV to MF | 67 | F | JAK2<sup>V617F</sup> | 0.69 | 3+/3 | 92.63% | Patient Blood |
| **MF6\*** | PMF | 72 | M | MPL | 0.94 | 3+/3 | 90.38% | Patient Blood |
| **CXCL8 non-secretor** |  |  |  |  |  |  |  |  |
| **MF10** | ET to MF | 67 | F | MPL | 0.24 | 1-2/+3 to 3+/3 | 18.0% | Patient Blood |
| **MF11** | PV to MF | 75 | M | JAK2<sup>V617F</sup> | 0.42 | 2-3+/3 | 6.98% | Patient Blood |
| **PV4\*** | PV | 70 | M | JAK2<sup>V617F</sup> | N/A | 1+, focal 2+/3 | 3.30% | Patient Blood |
| **ET2\*** | ET | 40 | M | CALR | 0.9 | 0-1+/3 | 3.71% | Patient Blood |
| **ET6** | ET | 73 | M | MPL | 0.24 | 1+/3 | 0.85% | Patient Blood |

MF: primary myelofibrosis; PV: polycythemia vera; ET: essential thrombocythemia. VAF: variant allele frequency
\*Denotes patients on whom integrated RNA-Seq/ATAC-Seq analysis was performed

**Supplemental Figure 4:**
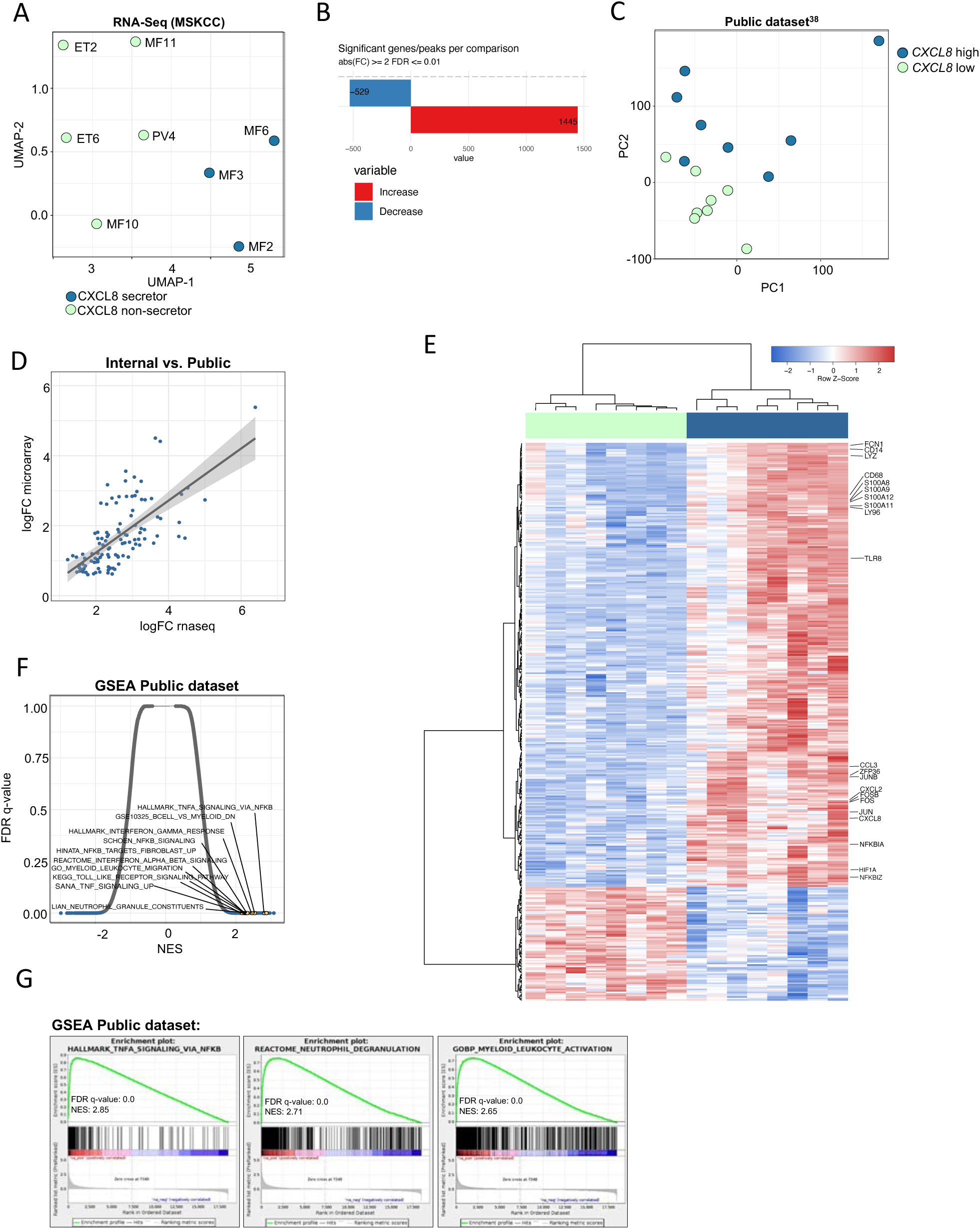
Extended RNA-Seq/ATAC-Seq patient data and publicly available MF gene expression dataset. **(A)** Uniform manifold approximation and projection (UMAP) visualization of N=9 MPN patient samples (see Supp. Table S2) available for gene expression (RNA-Seq) analysis. Each dot represents an individual patient. (Light blue represents CXCL8 “non-secretors;” dark blue represents CXCL8 “secretors” as identified by their single-cell cytokine secretion analysis). **(B)** Total number of differentially regulated genes (blue represents significantly down-regulated genes; red significantly up-regulated genes) of CXCL8 secretor vs. non-secretor MPN patients by a fold change cutoff of ± 2 and a FDR of 1%. **(C)** PCA plot demonstrating clustering of myelofibrosis patients segregated by CXCL8 expression levels and stratified by <20% and >80% from a publicly available microarray gene expression data set.^38^ Each dot represents an individual patient. Dark blue represents “*CXCL8* high” (>80%) MF patients; light blue represents “*CXCL8* low” (<20%) MF patients as based on their relative *CXCL8* expression level. **(D)** Correlation analysis comparing the internal (MSKCC) RNA-Seq dataset to the publicly-available MF gene expression data set (as per Norfo et al).^38^ **(E)** Hierarchal clustering demonstrating differentially expressed genes in select MF patients stratified by *CXCL8* expression level (light blue = *CXCL8* low; dark blue = *CXCL8* high). Blue, negative values; red, positive values. **(F)** Gene Set Enrichment Analysis (GSEA) demonstrating enriched pathways in “*CXCL8* high” vs. “*CXCL8* low” MF patients from the publicly-available dataset presented as normalized enrichment score (NES) by FDR q-value. **(G)** Representative GSEA plots demonstrating positive enrichment in pro-inflammatory signaling (TNFα via NF-κB) and myeloid cell/neutrophil activation gene sets in “*CXCL8* high” vs. “*CXCL8* low” MF patients from the publicly-available dataset.

**Supplemental Figure 5:**
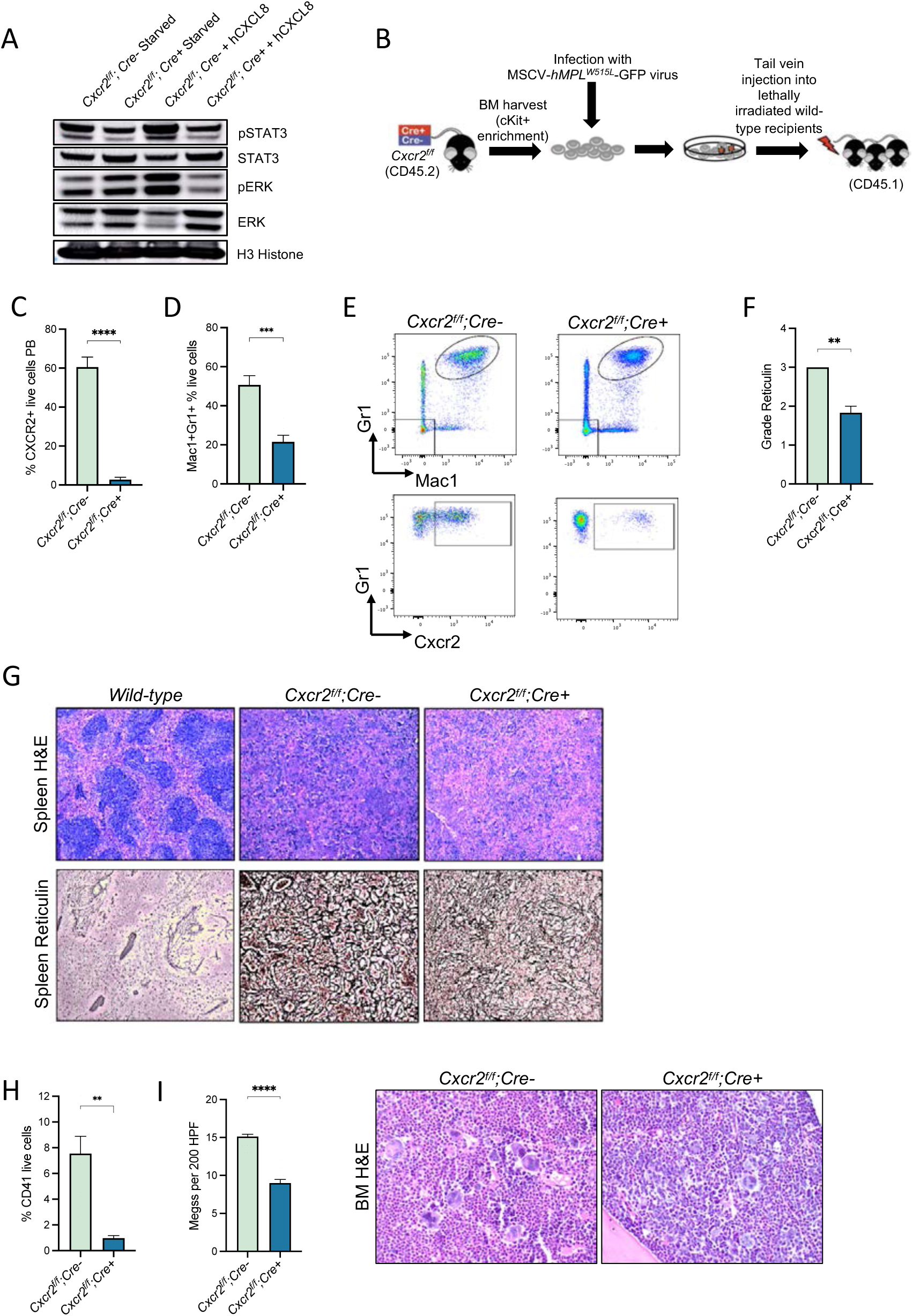
*Cxcr2^-/-^* knock-out validation, experimental set-up, and extended analysis of *Cxcr2^-/-^* hMPL^W515L^ mice. **(A)** Western blot analysis of canonical downstream Cxcr2 mediators in harvested murine bone marrow after 2 hours cultured with or without exogenous human CXCL8 for both *Cxcr2^f/f^;Cre^-^* wild-type (WT) and *Cxcr2^f/f^;Cre^+^* knock-out (KO) cells. **(B)** Schematic of experimental set up of the *Cxcr2^-/-^* KO hMPL^W515L^ transplant studies. **(C)** Confirmation of Cxcr2 loss on the surface of murine peripheral blood Mac1+Gr1+ cells from *Cxcr2^f/f^;Cre^+^* primary KO mice. ****p<0.0001. Data shown represent mean ± SEM. **(D)** Percent Mac1+Gr1+ mature neutrophil populations in peripheral blood of *Cxcr2^f/f^;Cre^+^* vs. *Cxcr2^f/f^;Cre^-^* hMPL^W515L^ transplanted mice at time of sacrifice. ***p<0.001. Data shown represent mean ± SEM. **(E)** Representative flow cytometry plots demonstrating reduction in peripheral blood Mac1+Gr1+ mature neutrophil populations and corresponding detectable Cxcr2 surface expression of *Cxcr2^f/f^;Cre^+^* vs. *Cxcr2^f/f^;Cre^-^* hMPL^W515L^ transplanted mice. **(F)** Bone marrow grade reticulin scores of *Cxcr2^f/f^;Cre^+^* vs. *Cxcr2^f/f^;Cre^-^* hMPL^W515L^ transplanted mice. **p<0.01. Data shown represent mean ± SEM. **(G)** Representative hematoxylin and eosin (H&E) and reticulin images of spleen sections from *Cxcr2^f/f^;Cre^+^* KO vs. *Cxcr2^f/f^;Cre^-^* WT hMPL^W515L^ mice. 20X magnification. **(H)** Percentage of CD41+ live cells as detected by flow cytometry in bone marrow of *Cxcr2^f/f^;Cre^+^* vs. *Cxcr2^f/f^;Cre^-^* hMPL^W515L^ transplanted mice at time of sacrifice. **p<0.01. Data shown represent mean ± SEM. **(I)** Recorded bone marrow megakaryocyte number (MKs) per high powered field (HPF) observed (left panel) and representative H&E stains of bone marrow (right panel) from *Cxcr2^f/f^;Cre^+^* KO vs. *Cxcr2^f/f^;Cre^-^* WT hMPL^W515L^ transplanted mice. N=4-7/arm. ****p<0.0001. 40X magnification. Data shown represent mean ± SEM.

**Supplemental Figure 6:**
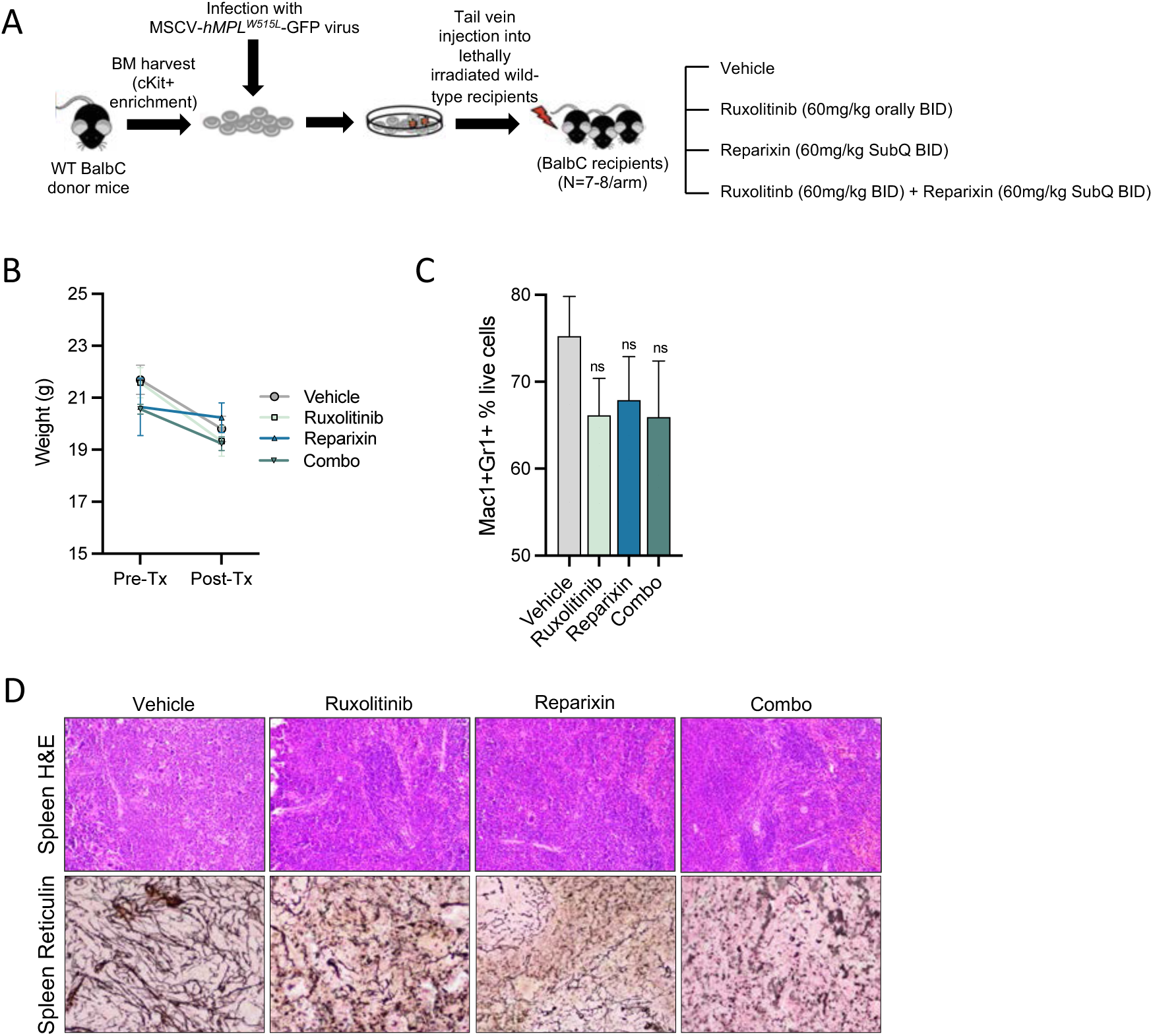
Pharmacologic inhibition of CXCR1/2 improves hematologic parameters and reticulin fibrosis in the hMPL^W515L^ adoptive transfer model of myelofibrosis. **(A)** Schematic of experimental outline for the reparixin/ruxolitinib *in vivo* drug trials. **(B)** Weights observed across treatment arms comparing the beginning and end of the 3-week trial period. N=6/arm. **(C)** Percentage of Mac1+Gr1+ neutrophil fractions in peripheral blood by flow cytometry of treated mice in comparison to vehicle at trial end point. ns=non-significant. Data shown represent mean ± SEM. **(D)** Representative H&E and reticulin images of hMPL^W515L^- diseased spleen treated with ruxolitinib, reparixin, or combination therapy compared with vehicle-treated hMPL^W515L^ mice. 20X magnification.

**Supplemental Figure 7:**
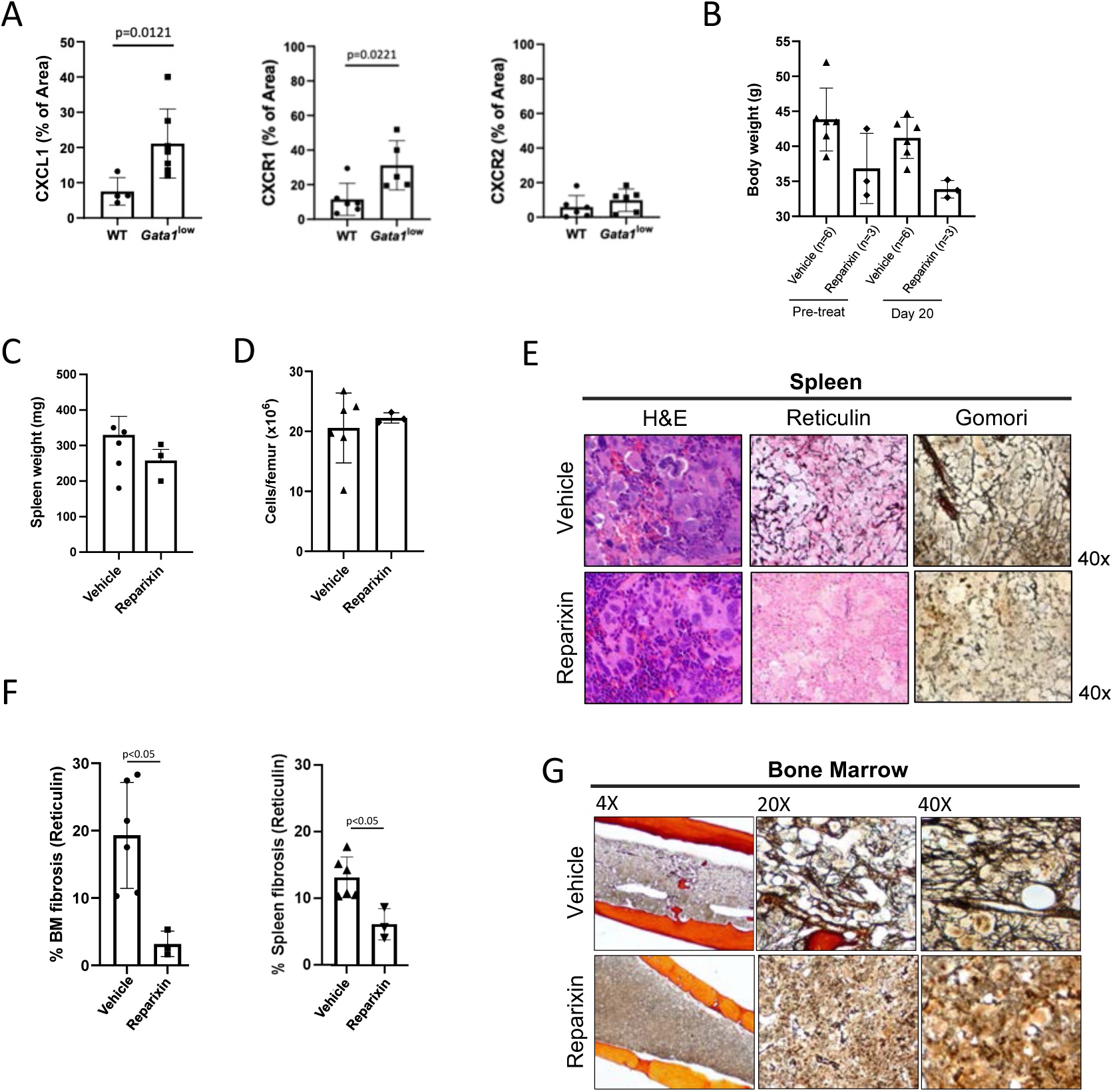
CXCR1/2 inhibition demonstrates therapeutic efficacy in the *Gata1^low^* model of myelofibrosis *in vivo*. **(A)** Quantification of Cxcl1, Cxcr1, and Cxcr2 abundance in *Gata1^low^* bone marrow compared to wild-type (WT) as quantified by % of area stained above threshold (see STAR methods). Data shown represent mean ± SD. **(B)** Body weights of trial mice prior to trial start and at the end of the trial period (day 20) for vehicle and reparixin cohorts. **(C)** Spleen weights of *Gata1^low^* mice in response to reparixin therapy compared to vehicle treated mice. Data shown represent mean ± SD. **(D)** Total bone marrow cellularity per femur of treated *Gata1^low^* mice following the 20 day *in vivo* trial period compared to vehicle treated mice. Data shown represent mean ± SD. **(E)** Representative images of H&E, reticulin, and Gomori stains of sectioned spleen samples of *Gata1^low^* mice following 20 days of reparixin therapy in comparison to vehicle treated mice. Magnification as indicated. **(F)** Quantification of reticulin in both bone marrow (BM; left panel) and spleen (right panel) of *Gata1^low^* mice in response to reparixin in comparison to vehicle treatment. Data shown represent mean ± SD. **(G)** Representative Gomori stains demonstrating degree fibrosis of sectioned bone marrow in reparixin vs. vehicle treated *Gata1^low^* mice. Magnification as indicated.

**Supplemental Figure 8:**
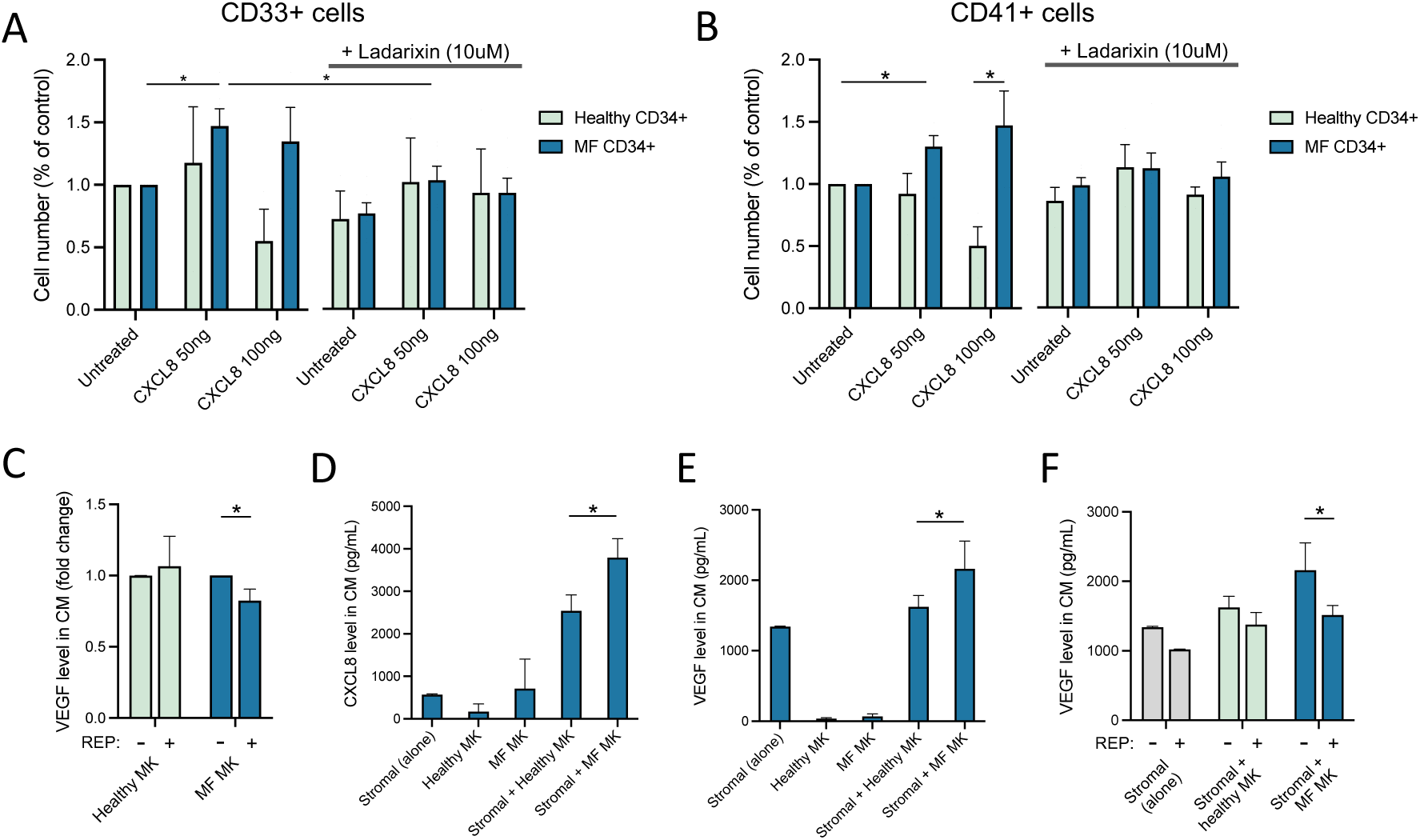
CXCR1/2 inhibition demonstrates efficacy against primary MPN cells *in vitro*. **(A)** Ratio of total CD33+ cell output relative to untreated of cultured healthy donor (HD; light blue) vs. myelofibrosis (MF; dark blue) CD34+ cells in response to exogenous CXCL8 doses (50ng vs. 100ng) with or without the CXCR1/2 inhibitor Ladarixin (10uM). *p<0.05. Data shown represent mean ± SD. **(B)** Ratio of total CD41+ cell output relative to untreated of cultured healthy donor (HD; light blue) vs. myelofibrosis (MF; dark blue) CD34+ cells in response to exogenous CXCL8 doses (50ng vs. 100ng) with or without the CXCR1/2 inhibitor Ladarixin (10uM). *p<0.05. Data shown represent mean ± SD. **(C)** Fold-change in detectable VEGF levels in conditioned media (CM) elicited by either healthy donor vs. MF megakaryocytes (MKs) with or without the addition of reparixin (REP; 10uM). *p-value <0.05. Data shown represent mean ± SD. **(D)** Total CXCL8 levels detected in conditioned media of various co-culture conditions (stromal cells alone +/- healthy donor or MF megakaryocytes [MK]). *p-value <0.05. Data shown represent mean ± SD. **(E)** Total VEGF levels detected in conditioned media with various co-culture conditions (stromal cells alone +/- healthy donor vs. MF MKs). *p-value <0.05. Data shown represent mean ± SD. **(F)** Total levels of VEGF in conditioned media (CM) of cultured stromal cells, either alone or together with healthy vs. MF MKs with or without the addition of reparixin (REP; 10uM). *p-value <0.05. Data shown represent mean ± SD.

**Supplemental Table 4:** Selected myelofibrosis (MF) patient samples for *in vitro* studies.

| ID | Diagnosis | Age | Sex | Driver Mutation | Grade Reticulin Fibrosis | Experiments performed |
| --- | --- | --- | --- | --- | --- | --- |
| MF1 | PMF | 75 | M | JAK2<sup>V617F</sup> | Not reported | CFA, JAK2 genotyping, CXCR1/2 test |
| MF2 | PMF | 88 | F | JAK2<sup>V617F</sup> | 3 | CFA, JAK2 genotyping, CXCR1/2 test , qPCR |
| MF3 | PET-MF | 57 | F | JAK2<sup>V617F</sup> | Not reported | CFA, JAK2 genotyping, CXCR1/2 test , co-culture |
| MF4 | PET-MF | 75 | F | JAK2<sup>V617F</sup> | Not reported | CFA, JAK2 genotyping, CXCR1/2 test, proliferation |
| MF5 | PMF | 51 | M | CALR | 3 | CFA,CXCR1/2 test , qPCR |
| MF6 | PMF | 74 | M | JAK2<sup>V617F</sup> | 2 | CFA, JAK2 genotyping, CXCR1/2 test , proliferation, co-culture, qPCR |
| MF7 | PMF | 74 | F | CALR | 3 | CFA, CXCR1/2 test , qPCR |
| MF8 | PET-MF | 57 | M | CALR | 2-3 | CFA, CXCR1/2 test |
| MF9 | PET-MF | 81 | F | JAK2<sup>V617F</sup> | 2 | CFA, qPCR |
| MF10 | PET-MF | 47 | F | CALR | 3 | CFA, qPCR |
| MF11 | PMF | 74 | M | JAK2<sup>V617F</sup> | Not reported | CFA+JAK2genotyping |
| MF12 | PPV-MF | 75 | M | MPL | 3 | CFA |
| MF13 | PET-MF | 75 | F | JAK2<sup>V617F</sup> | Not reported | proliferation |
| MF14 | PET-MF | 63 | F | CALR | 1-2 | CFA, proliferation |
| MF15 | PET-MF | 77 | F | CALR | Not reported | CFA, proliferation, Co-culture |
| MF16 | PMF/MPN-BP | 63 | F | CALR | Not reported | proliferation |
| MF17 | PMF | 65 | M | JAK2<sup>V617F</sup> | 3 | CXCR1/2 test |
| MF18 | MF |  |  | JAK2<sup>V617F</sup> | Not reported | CXCR1/2 test |
| MF19 | PPV-MF | 65 | F | JAK2<sup>V617F</sup> | 3 | CXCR1/2 test |
| MF20 | PPV-MF | 64 | F | JAK2<sup>V617F</sup> | Not reported | CXCR1/2 test |
| MF21 | PMF | 73 | M | JAK2<sup>V617F</sup> | 3 | CXCR1/2 test |
| MF22 | PET-MF | 52 | F | CALR | 3 | CXCR1/2 test |
| MF23 | PMF | 62 | M | JAK2<sup>V617F</sup> | 2-3 | CXCR1/2 test |

PMF: primary myelofibrosis; MF: myelofibrosis; PET-MF: post-essential thrombocythemia myelofibrosis; PPV-MF: post-polycythemia vera myelofibrosis; CFA: Colony formation assay.

